# Benchmarking of computational error-correction methods for next-generation sequencing data

**DOI:** 10.1101/642843

**Authors:** Keith Mitchell, Jaqueline J. Brito, Igor Mandric, Qiaozhen Wu, Sergey Knyazev, Sei Chang, Lana S. Martin, Aaron Karlsberg, Ekaterina Gerasimov, Russell Littman, Brian L. Hill, Nicholas C. Wu, Harry Yang, Kevin Hsieh, Linus Chen, Eli Littman, Taylor Shabani, German Enik, Douglas Yao, Ren Sun, Jan Schroeder, Eleazar Eskin, Alex Zelikovsky, Pavel Skums, Mihai Pop, Serghei Mangul

**Author notes:** These authors contributed equally to the paper (Keith Mitchell, Jaqueline J. Brito, Igor Mandric).

## Abstract

**Background:** Recent advancements in next-generation sequencing have rapidly improved our ability to study genomic material at an unprecedented scale. Despite substantial improvements in sequencing technologies, errors present in the data still risk confounding downstream analysis and limiting the applicability of sequencing technologies in clinical tools. Computational error-correction promises to eliminate sequencing errors, but the relative accuracy of error correction algorithms remains unknown.

**Results:** In this paper, we evaluate the ability of error-correction algorithms to fix errors across different types of datasets that contain various levels of heterogeneity. We highlight the advantages and limitations of computational error correction techniques across different domains of biology, including immunogenomics and virology. To demonstrate the efficacy of our technique, we apply the UMI-based high-fidelity sequencing protocol to eliminate sequencing errors from both simulated data and the raw reads. We then perform a realistic evaluation of error correction methods.

**Conclusions:** In terms of accuracy, we find that method performance varies substantially across different types of datasets with no single method performing best on all types of examined data. Finally, we also identify the techniques that offer a good balance between precision and sensitivity

## Introduction

Rapid advancements in next-generation sequencing have improved our ability to study the genomic material of a biological sample at an unprecedented scale and promise to revolutionize our understanding of living systems^1^. Sequencing technologies are now the technique of choice for many research applications in human genetics, immunology, and virology^1,2^. Modern sequencing technologies dissect the input genomic DNA (or reverse transcribed RNA) into millions of nucleotide sequences, which are known as reads. Despite constant improvements in sequencing technologies, the data produced by these techniques remain biased by the introduction of random and systematic errors. Sequencing errors typically occur in approximately 0.1-1% of bases sequenced; such errors are more common in reads with poor-quality bases where sequencers misinterpret the signal or when the wrong nucleotide is incorporated. Errors are introduced at the sequencing step via incorporation of faults and even occur in reads with few poor-quality bases per read^3^. Additional errors, such as polymerase bias and incorporation errors, may be introduced during sample preparation, amplification, or library preparation stages^4^. Data containing sequencing errors limit the applicability of sequencing technologies in clinical settings^5^. Further, the error rates vary across platforms^6^; the most popular Illumina-based protocols can produce approximately one error in every one thousand nucleotides^7^.

In order to better understand the nature of and potential solutions for sequencing errors, we conducted a comprehensive benchmarking study of currently available error correction methods. We identified numerous effects that various sequencing settings, and the different parameters of error correction methods, can have on the accuracy of output from error correction methods. We also investigated the advantages and limitations of computational error correction techniques across different domains of biology, including immunogenomics and virology.

Computational error-correction techniques promise to eliminate sequencing errors and improve the results of downstream analyses (**Figure 1a**)^8^. Many computational error correction methods have been developed to meet growing demand for accurate sequencing data in the biomedical community ^9–11^. Despite the availability of many error correction tools, thoroughly and accurately eliminating errors from sequencing data remains a challenge. First, currently available molecular-based techniques for correcting errors in sequencing data (e.g., ECC-Seq^12^) usually carry an increased computational cost which limits scalability across a large number of samples. Second, our lack of a systematic comparison of error correction methods impedes the optimal integration of these tools into standardized next-generation sequencing data analysis pipelines.

**Figure 1.**
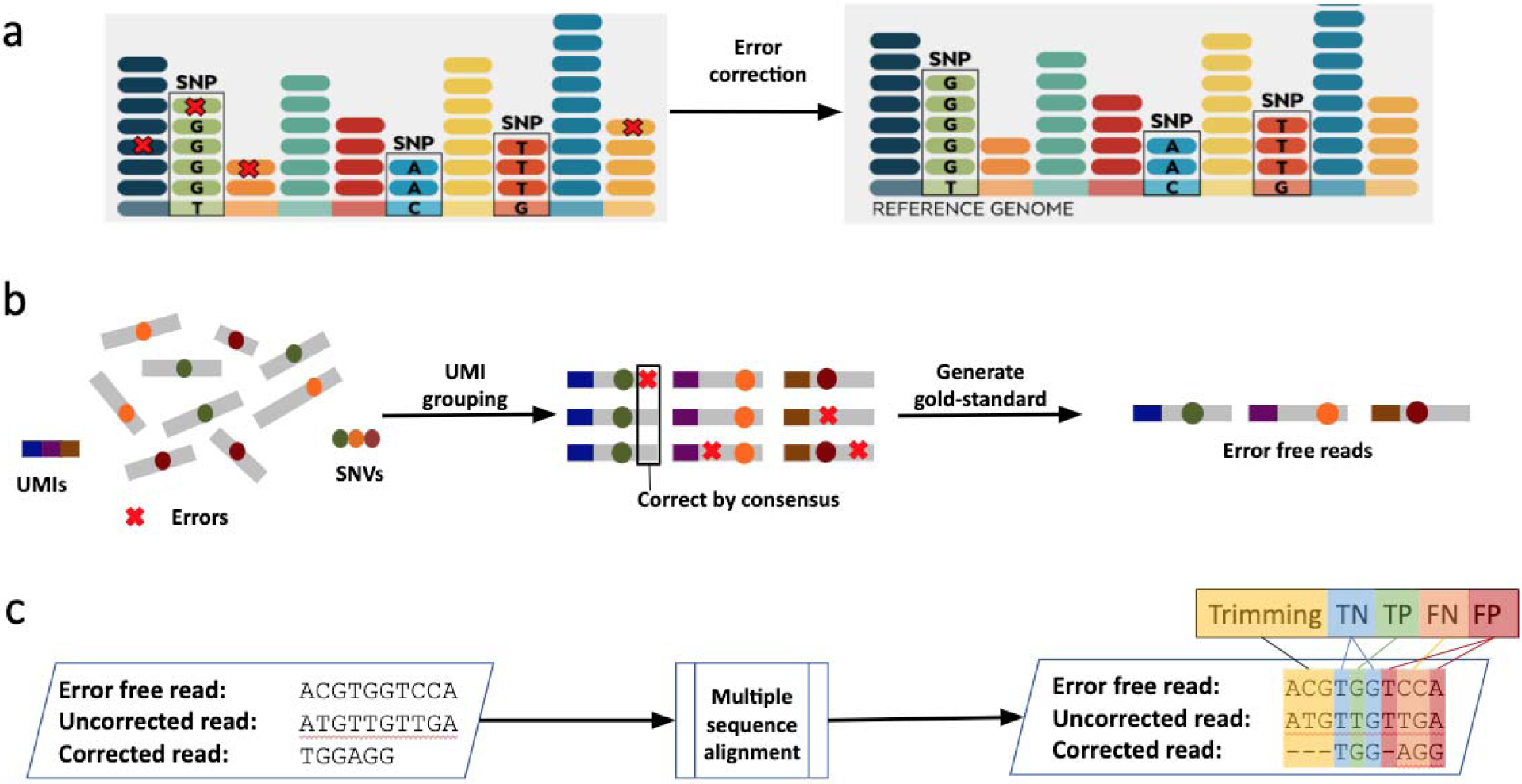
Study design for benchmarking computational error correction methods. **(a)** Schematic representation of the goal of error correction algorithms. Error correction aims to fix sequencing errors while maintaining the data heterogeneity. **(b)** Error-free reads for gold standard were generated using UMI-based clustering. Reads were grouped based on matching UMIs and corrected by consensus, where an 80% majority was required to correct sequencing errors without affecting naturally occurring Single Nucleotide Variations (SNVs). **(c)** Framework for evaluating the accuracy of error correction methods. Multiple sequence alignment between the error-free, uncorrected (original) and corrected reads was performed to classify bases in the corrected read. Bases fall into the category of Trimming, True Negative (TN), True Positive (TP), False Negative (FN), and False Positive (FP).

Previous benchmarking studies^13,14^ lacked a comprehensive experimental gold standard^15^; instead, these early benchmarking efforts relied on simulated data and real reads which were uniquely aligned to the reference genome. In addition, error correction algorithms have undergone significant development since the earlier benchmarking studies, and the performance of the newest methods has not yet been evaluated. Other studies^16^ provide a detailed description of available error correction tools yet lack the benchmarking results. The efficiency of today’s error correction algorithms, when applied to the extremely heterogeneous populations composed of highly similar yet distinct genomic variants, is presently unknown. The human immune repertoire, a collection of diverse B and T cell receptor clonotypes, is an excellent example of a heterogeneous population with need for reliable error correction. The increased heterogeneity of such datasets and the presence of low-frequency variants further challenges the ability of error-correction methods to fix sequencing errors in the data.

In this paper, we evaluate the ability of error-correction algorithms to fix errors across different types of datasets with various levels of heterogeneity. In doing so, we produce a gold standard that provides an accurate baseline for performing a realistic evaluation of error correction methods. We highlight the advantages and limitations of computational error correction techniques across different domains of biology, including immunogenomics and virology. For example, we challenged the error correction methods with data derived from diverse populations of T cell receptor clonotypes and intra-host viral populations. To define a gold standard for error correcting methods, we applied a Unique Molecular Identifier (UMI)-based high-fidelity sequencing protocol (also known as safe-SeqS)^17,18^ and eliminated sequencing errors from raw reads.

## Results

### Gold standard datasets

We used both simulated and experimental gold standard datasets derived from human genomic DNA, human T cell receptor repertoires, and intra-host viral populations. The datasets we used correspond to different levels of heterogeneity. The difficulty of error correction increases as the dataset becomes more heterogeneous. The least heterogeneous datasets were derived from human genomic DNA (**D1 dataset**) (**Table 1**). The most heterogeneous datasets were derived from the T cell receptor repertoire and from a complex community of closely related viral mutant variants (known as quasispecies).

**Table 1:** Overview of the gold standard datasets.

|  | D1 | D2 | D3 | D4 | D5 |
| --- | --- | --- | --- | --- | --- |
| Technology | Whole genome sequencing | T cell receptor sequencing | T cell receptor sequencing | Viral sequencing | Viral sequencing |
| Heterogeneity | Low | High | High | High | High |
| Technique | Simulated | UMI-based error-free reads | Simulated | UMI-based error-free reads | Haplotype-based error-free reads |
| Number of samples | 12 | 8 | 6 | 1 | 11 |

To generate error-free reads for the D2 and D4 datasets, we used a UMI-based high-fidelity sequencing protocol (also known as safe-SeqS)^17,18^, which is capable of eliminating sequencing errors from the data (**Figure 1b**). A high-fidelity sequencing protocol attaches the UMI to the fragment prior to amplification of DNA fragments. After sequencing, the reads that originated from the same biological segment are grouped into clusters based on their UMI tags. Next, we applied an error-correction procedure inside each cluster of biological segments. In cases where at least one nucleotide inside the UMI cluster lacks the support of 80% of reads, we were not able to generate consensus error-free read; in other words, if 80% have the reads have the same nucleotide we consider that nucleotide a correct one. When the nucleotide lacks support of 80% of reads, all reads from these UMI clusters were disregarded (**Figure 1c**). We used UMI-based clustering to produce error-free reads for the D2 and D4 datasets. Both the D1 and D3 datasets were produced by computational simulations using a customized version of the tool WgSim^19^ **(Additional file 1: Fig. S1)**.

We applied a haplotype-based error correction protocol to eliminate sequencing errors from the D5 dataset, composed of five HIV-1 subtype B haplotypes that were mixed *in-vitro*^*20*^. First, we determined the haplotype of origin for each read by aligning reads on the set of known haplotypes obtained from the mixture. Sequencing errors were corrected by replacing bases from reads, with the bases from the haplotype of origin. We varied the number of haplotypes and the similarity of haplotypes present in the HIV-1 mixture. In addition, we varied the rate of sequencing errors in the data.

Availability of both error-free reads and the original raw reads carrying errors provide an accurate, robust baseline for performing a realistic evaluation of error correction methods. For our benchmarking study, we examined both experimental data and simulated data. Simulated data contain reads with various lengths and coverage rates to estimate the effect of such sequencing parameters on the accuracy of error correction. A detailed description of the dataset used and the corresponding protocol to prepare gold standard dataset is provided in the **Methods section**.

### Choice of error correction methods

We chose the most commonly-used error correction tools to assess the ability of current methods to correct sequencing errors. The following algorithms were included in our benchmarking study: Coral^21^, Bless^10^, Fiona^10,22^, Pollux^11^, BFC^23^, Lighter^24^, Musket^9^, Racer^25^, RECKONER^24,26^, and SGA^27^. We excluded HiTEC and replaced it with Racer, as was recommended by the developers of HiTEC. We also excluded tools solely designed for non-Illumina based technologies^28^ and tools which are no longer supported. We summarized the details of each tool, including the underlying algorithms and the data structure (**Table 2)**. To assess the simplicity of the installation process for each method, we describe the software dependencies in **Table 3**. Commands required to run each of the tools are available in the Supplementary Materials **(Additional file 1: Table S1)** and at https://github.com/Mangul-Lab-USC/benchmarking.error.correction. The data preparation for evaluation is described at the Supplementary Materials (**Additional file 1: Supplemental Note 2**).

**Table 2:**
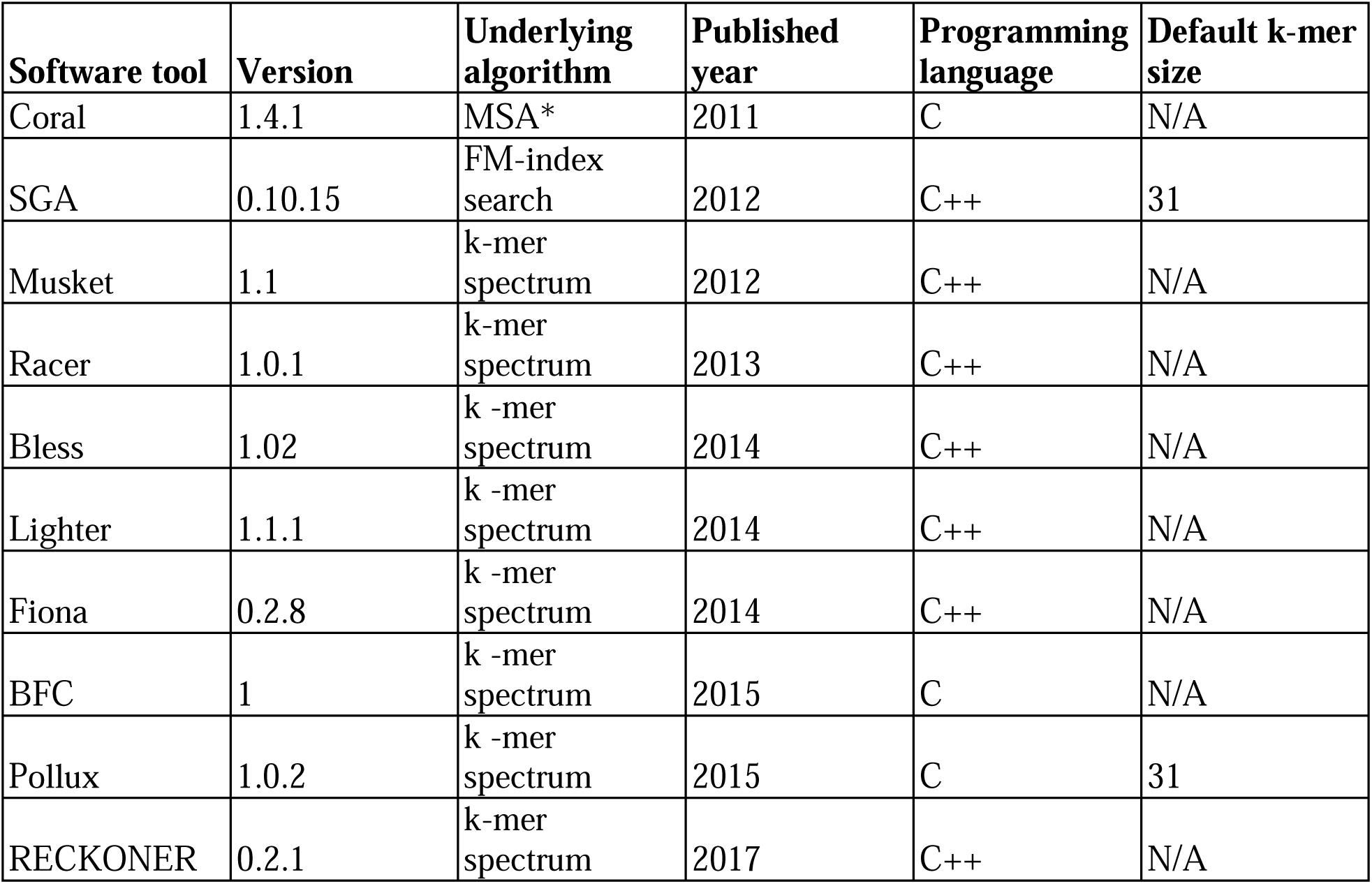
Summary of error correction methods parameters and publication details. Error correction methods are sorted by the year of publication (indicated in column “Published Year”). We documented underlying algorithm (indicated in column “Underlying algorithm”), version of the error correction tool used (indicated in column “Version”), and the name of the software tool (indicated in column “Software tool”).

| **Software tool** | **Version** | **Underlying algorithm** | **Published year** | **Programming language** | **Default k-mer size** |
| --- | --- | --- | --- | --- | --- |
| Coral | 1.4.1 | MSA* | 2011 | C | N/A |
| SGA | 0.10.15 | FM-index search | 2012 | C++ | 31 |
| Musket | 1.1 | k-mer spectrum | 2012 | C++ | N/A |
| Racer | 1.0.1 | k-mer spectrum | 2013 | C++ | N/A |
| Bless | 1.02 | k-mer spectrum | 2014 | C++ | N/A |
| Lighter | 1.1.1 | k-mer spectrum | 2014 | C++ | N/A |
| Fiona | 0.2.8 | k-mer spectrum | 2014 | C++ | N/A |
| BFC | 1 | k-mer spectrum | 2015 | C | N/A |
| Pollux | 1.0.2 | k-mer spectrum | 2015 | C | 31 |
| RECKONER | 0.2.1 | k-mer spectrum | 2017 | C++ | N/A |

**Table 3:**
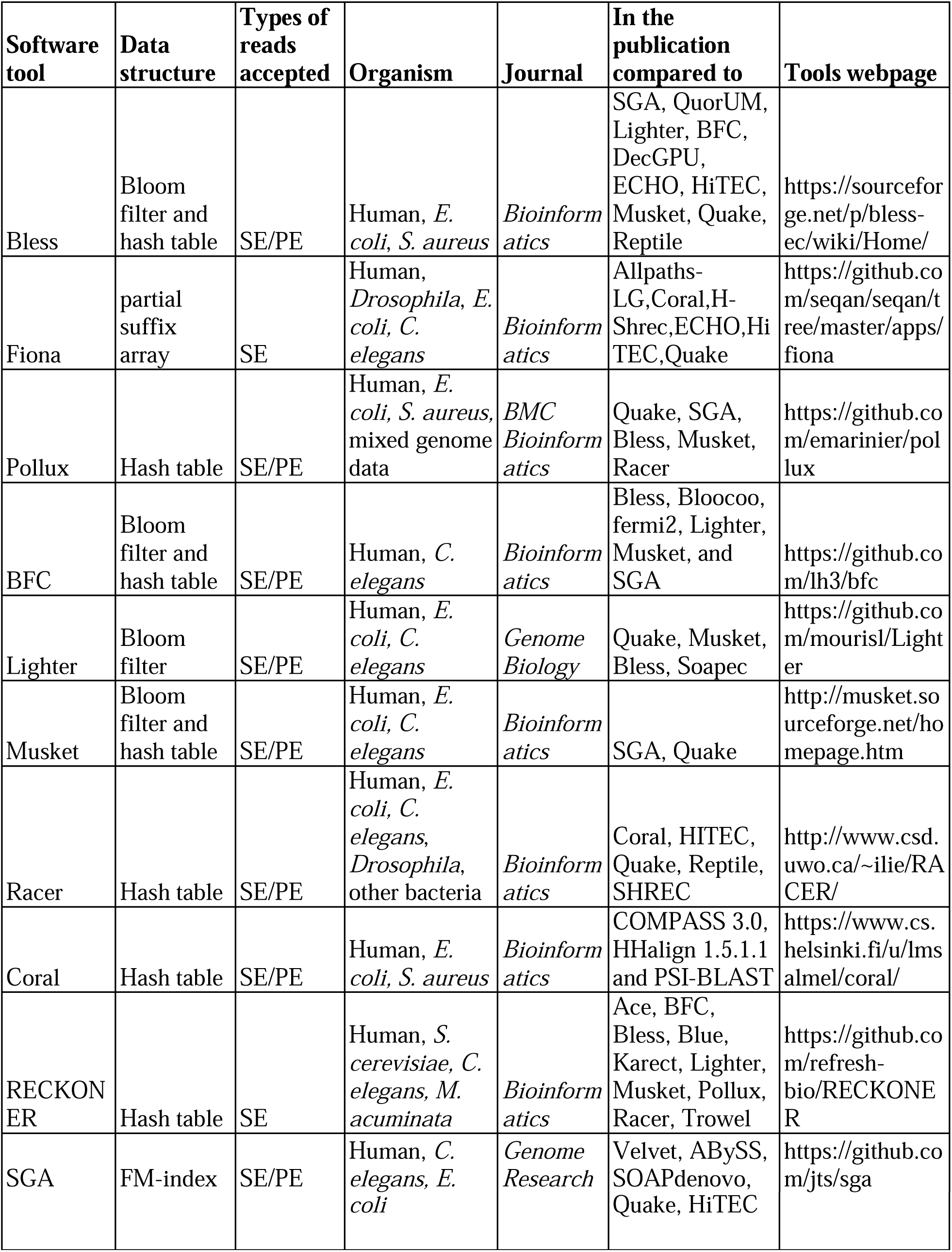
Summary of technical characteristics of the error correction methods assessed in this study.

### Evaluation of the accuracy and performance of error correction methods

We used an extensive set of evaluation metrics to assess the accuracy and performance of each error correction method. We defined true positives (TP) as errors that were correctly fixed by the error correction tool; false positives (FP) as correct bases that were erroneously changed by the tool; false negatives (FN) as erroneous bases not fixed or incorrectly fixed by the tool, and true negatives (TN) as correct bases which remain unaffected by the tool (**Figure 1b**) (**Additional file 1: Fig. S2**).

We used the gain metric^13^ to quantify the performance of each error correction tool. Positive gain represent an overall positive effect of the error correction algorithm, whereas a negative gain shows that the tool performed more incorrect actions then correct actions. A gain of 1.0 means the error correction tool made all necessary corrections without any FP alterations (**Table 3**). We defined precision as the proportion of proper corrections among the total number of corrections performed by the error correction tool. Sensitivity evaluates the proportion of fixed errors among all existing errors identified in the data; in other words, sensitivity indicates which algorithms are correcting the highest majority of induced errors ^29^. Finally, we checked if the error correction methods remove the bases in the beginning or the end of corrected reads. Removing the bases may correspond with an attempt to correct deletion (TP trimming) or may simply remove a correct base (FP trimming) (**Additional file 1: Fig. S3**).

**Table 3:** Evaluation of the accuracy of error correction methods. **(a)** Precision evaluates the proportion of proper corrections among the total number of performed corrections. INDEL refers to insertion/deletion polymorphism. **(b)** Sensitivity evaluates the proportion of fixed errors among all existing errors in the data. **(c)** Gain represents whether an algorithm is producing an overall benefit (more TP then FP) or is having a negative effect (more FP then TP). Values ranging from 1.0 to, but not including, 0.0 represent a benefit; 0.0 is neutral; and less than 0.0 is considered a negative effect. **(d)** Trim efficiency is the proportion of trimmed bases from the tool that were considered to be TP trimming. **(e)** Trim percent is the proportion of nucleotides trimmed out of all nucleotides analyzed.

|  | Metric Name | Metric Formula |
| --- | --- | --- |
| **a.** | Precision | $= \begin{cases} \frac{TP}{TP + FP + FP_{INDEL}}, & TP + FP + FP_{INDEL} > 0 \\ 0, & TP + FP + FP_{INDEL} = 0 \end{cases}$ |
| **b.** | Sensitivity | $= \begin{cases} \frac{TP}{TP + FN}, & TP + FN > 0 \\ 0, & TP + FN = 0 \end{cases}$ |
| **c.** | Gain | $= \begin{cases} \frac{TP - (FP + FP_{INDEL})}{TP + FN}, & TP + FN > 0 \\ 0, & TP + FN = 0 \end{cases}$ |
| **d.** | Trim Percent | $= \begin{cases} \frac{TP_{TRIM}}{TotalBases}, & TotalBases > 0 \\ 0, & TotalBases = 0 \end{cases}$ |
| e. | Trim Efficiency | $= \begin{cases} \frac{TP_{TRIM}}{TP_{TRIM} + FP_{TRIM}}, & TP_{TRIM} + FP_{TRIM} > 0 \\ 0, & TP_{TRIM} + FP_{TRIM} = 0 \end{cases}$ |

### Correcting errors in the whole genome sequencing data

We evaluated the efficacy of currently available error correction methods in fixing errors introduced to whole genome sequencing (WGS) reads using various coverage settings **(D1 dataset)** (**Table 1**). First, we explored the effect of k-mer size on the accuracy of error correction methods. An increase in k-mer size typically offers increased accuracy of error correction. In some cases, increased k-mer size has no effect on the accuracy of error correction (**Additional file 1: Fig. S4a-f**). We used the best k-mer size for all surveyed methods (**Additional file 1: Table S2**). The Lighter method for WGS human data with 32x coverage performs best with k-mer size of 30bp (**Additional file 1: Fig. S4f**). For other coverages, Lighter usually performs best with k-mer size of 20bp, which was chosen in those cases. Overall, the increase in k-mer size results in a decreased number of corrections for all the tools with regards to WGS data (**Additional file 1: Fig. S5**).

Our results show that Pollux and Musket make the largest number of corrections across all coverage settings when applied to the D1 dataset with coverage of 4x or less (**Additional file 1: Fig. S5**). In general, higher coverage allows error correction methods to make more corrections and fix more errors in the data. For the majority of the surveyed methods higher coverage also results in decreasing the number of false corrections (**Additional file 1: Fig. S6**). For the vast majority of the tools (except Lighter and Racer) gain constantly increases with coverage increase (**Figure 2a**). For the majority of the error correction tools in our study, the gain becomes positive only for coverage of 4x or higher. The only methods that demonstrated positive gain for 2x coverage were SGA and Coral. For coverage of 1x, Coral was the only method able to maintain a positive gain (**Figure 2a**). Coverage level also had a strong impact on both precision and sensitivity (**Figure 2b-c**). Except for Coral, none of the methods were able to correct more than 80% of the data for datasets with coverage of 2x or less (**Figure 2c**). Coverage of 32x allowed most methods to correct more than 95% of errors with high precision (**Figure 2f**). The error correction tools typically trimmed a minor portion of the reads. We compared the trimming rates and trends across the error correction methods. Overall, the majority of error correction tools trim a small percentage of bases. The only exception was Bless, which trimmed up to 29% of bases (**Additional file 1: Fig. S7**). The vast majority of trimmed bases were correct bases (**Additional file 1: Fig. S8-S9**).

**Figure 2.**
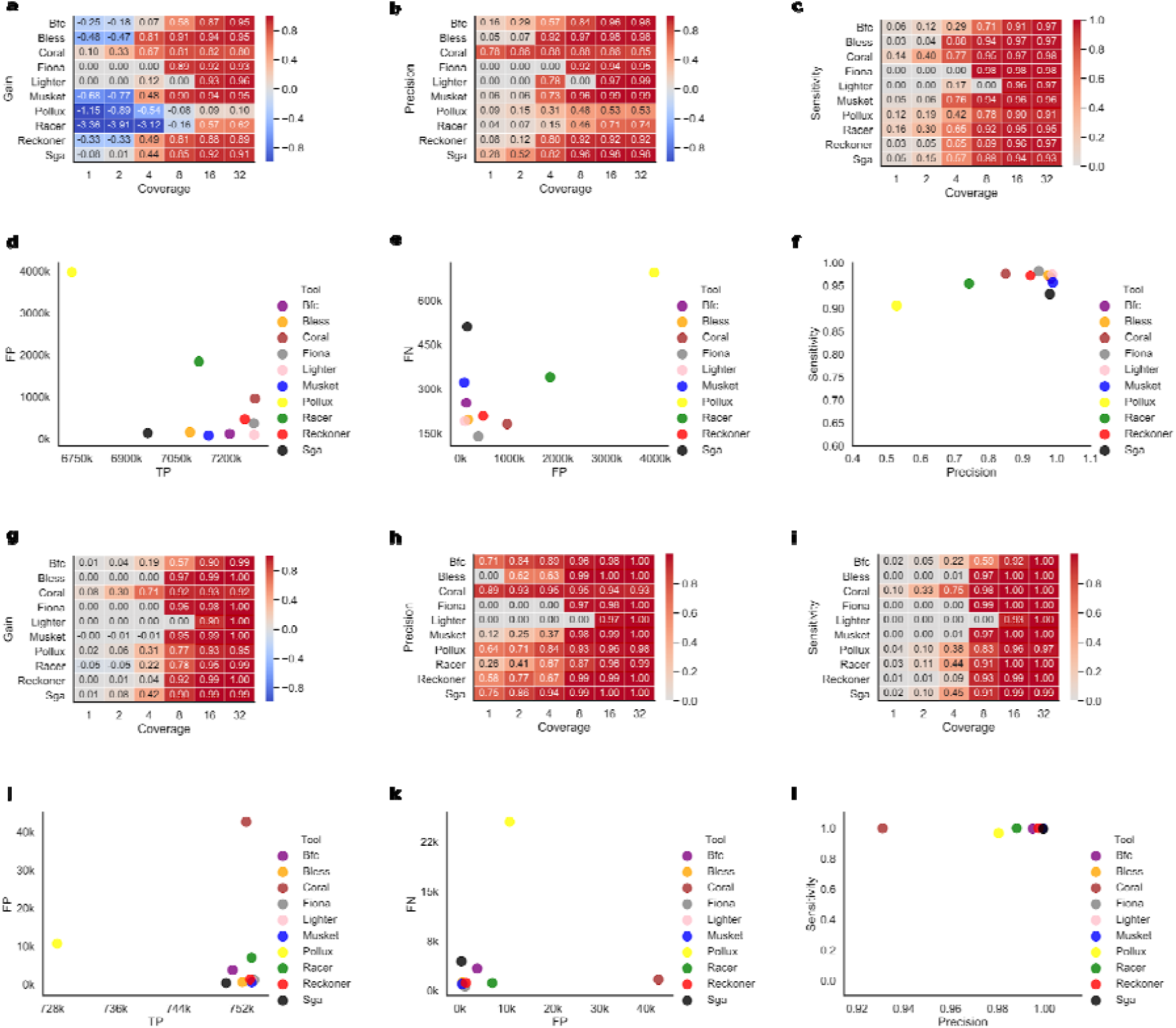
Correcting errors in whole genome sequencing data (D1 dataset). For each tool, the best k-mer size was selected. **(a-f)** WGS human data. **(g-l)** WGS *E. coli* data. **(a**,**g)** Heatmap depicting the gain across various coverage settings. Each row corresponds to an error correction tool, and each column corresponds to a dataset with a given coverage. **(b**,**h)** Heatmap depicting the precision across various coverage settings. Each row corresponds to an error correction tool, and each column corresponds to a dataset with a given coverage. **(c**,**i)** Heatmap depicting the sensitivity across various coverage settings. Each row corresponds to an error correction tool, and each column corresponds to a dataset with a given coverage. **(d**,**j)** Scatter plot depicting the number of TP corrections (x-axis) and FP corrections (y-axis) for datasets with 32x coverage. **(e**,**k)** Scatter plot depicting the number of FP corrections (x-axis) and FN corrections (y-axis) for datasets with 32x coverage. **(f**,**l)** Scatter plot depicting the sensitivity (x-axis) and precision (y-axis) for datasets with 32x coverage.

We have also compared the accuracy of error correction algorithm on *E. coli* WGS data. The relative performance of error correction methods was similar to the WGS human data. However, the differences in performance between the tools on *E. coli* data were smaller compared to human data specially for high coverage data (**Figure 2d-f,j-l**). Notably, many tools are able to maintain excellent performance (gain above 90%) even for coverages as low as 8x. The tool with the best performance for low coverage WGS when applied to both human and *E. coli* data was Coral, which was able to maintain positive gain even for 1x WGS data for both *E. coli* and human data (**Figure 2g**). Precision of error correction tools on *E. coli* was generally high even for low coverage data (**Figure 2h**). Many tools are able to achieve sensitivity above 90% even for 8x coverage (**Figure 2i**). Similar to human data, majority of the tools are able to maintain a good balance between precision and sensitivity for 32x WGS data (**Figure 2f**,**l**).

We have also investigated the performance of the tools in the low complexity regions. Excluding the low complexity regions results in a moderate improvement of accuracy for the majority of the tools. The largest difference in performance between low complexity regions and the rest of the genome was evident in results generated by Racer and Pollux. Notably, the only tool with a negative gain for low complexity regions was Pollux (**Additional file 1: Fig. S10**).

We have also compared CPU time and the maximum amount of RAM used by each of the tools based on WGS data (**Additional file 1: Fig. S11**). Bless, Racer, RECKONER, Lighter, and BFC were the fastest tools and were able to correct the errors in less than two hours for the WGS sample corresponding to chromosome 21 with 8x coverage. Other tools required more than five hours to process the same samples. The tools with lowest memory footprint were Lighter, SGA, and Musket, requiring less than 1GB of RAM to correct the reads in the samples. The tool with the highest memory footprint was Coral, requiring more than 9GB of RAM to correct the errors.

### Correcting errors in the TCR sequencing data

We compared the ability of error correction methods to fix the errors in reads derived from the T cell receptor (TCR) repertoire **(D2 and D3 datasets)** (**Table 1**). We investigated the effect of k-mer size using real TCR-Seq data derived from 8 individuals diagnosed with HIV (D2 dataset) and simulated TCR-Seq data (D3 dataset). For D2 dataset, error-free reads for a gold standard were generated by consensus using UMI-based clustering (see the **Methods section**). Similarly to our study of the WGS data, we explored the effect of k-mer size on the accuracy of error correction methods for TCR-Seq data. As we observed with the WGS data, with TCR-Seq data an increase in k-mer size improves the gain for some of the tools, while for other tools it has no effect (**Additional file 1: Fig. S12-S13**). We used the best k-mer size for all surveyed methods (**Additional file 1: Table S3-S4**).

We have used simulated TCR-Seq data (D3 dataset) to compare the performance of the error correction tools across various coverages. All error correction tools are able to maintain positive gain on simulated TCR-Seq data across various coverages (**Additional file 1: Fig. S14**). The vast majority of surveyed tools also maintain high precision rates (0.76-0.99) (**Additional file 1: Fig. S15**). We observed large variation in sensitivity across the tools and coverages. For several error correction methods, sensitivity drops when the coverage rate increases (**Additional file 1: Fig. S16**). Next, we have used real TCR-Seq data to compare the performance of error correction tools. The highest accuracy is achieved using the Lighter method, followed by Fiona and BFC (**Figure 3a**).

**Figure 3.**
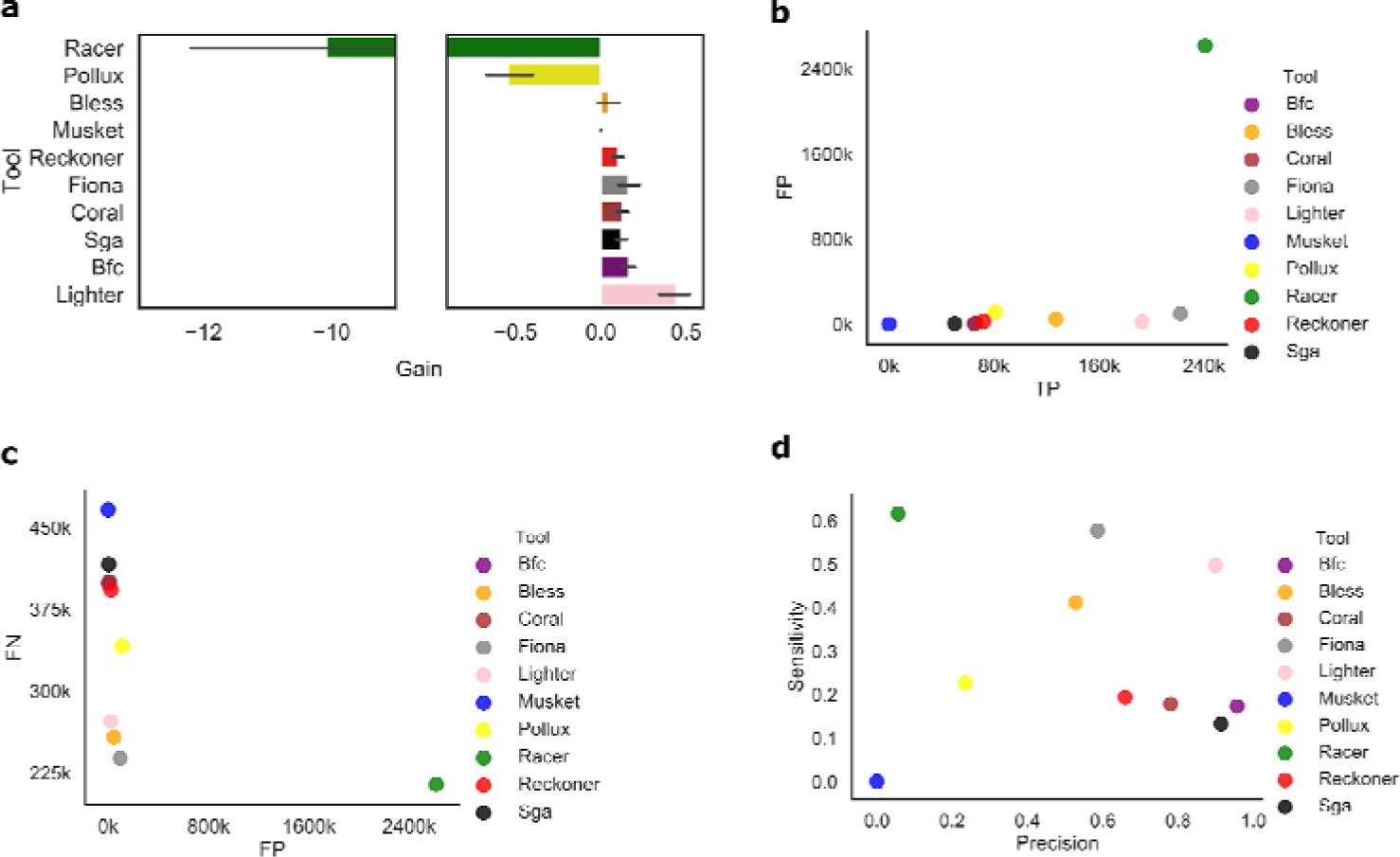
Correcting errors in TCR-Seq data (D2 dataset). For all plots, the mean value across 8 TCR-Seq samples is reported for each tool. **(a)** Bar plot depicting the gain across various error correction methods. **(b)** Scatter plot depicting the number of TP corrections (x-axis) and FP corrections (y-axis). **(c)** Scatter plot depicting the number of FP corrections (x-axis) and FN corrections (y-axis). **(d)** Scatter plot depicting the sensitivity (x-axis) and precision (y-axis) of each tool.

Lighter achieves a desirable balance between precision and sensitivity, and generally manifest similar performance according to all metrics, including number of TPs and FPs (**Figure 3b-d**). Due to the increased number of ignored errors (FNs), SGA demonstrates the lowest sensitivity among all error correction methods (**Figure 3d**). Similarly to WGS data, the majority of error correction tools do not trim or only trim a minor portion of the reads. Similar to results generated from WGS datasets, only a small number of reads were trimmed. Typically, the majority of trimmed bases were correct bases **(Additional file 1: Fig. S17)**.

### Correcting errors in the viral sequencing data

We compared the ability of error correction methods to fix the errors in reads derived from the heterogeneous viral populations (**D4 dataset**) (**Table 1**). First, we explored the effect of k-mer size on the accuracy of error correction methods for viral sequencing data **(Additional file 1: Fig. S18)**. For several error correction methods, k-mer size does not have substantial effect on accuracy of error correction. The best k-mer size was chosen for each tool (**Additional file 1: Table S5**). Majority of the methods are able to maintain precision above 80% (**Figure 4a**). Methods with the best balance between precision and sensitivity was Fiona, which also maintained the highest f-score (**Figure 4a**). None of the methods was able to correct more than 54% of errors (**Figure 4b**).

**Figure 4.**
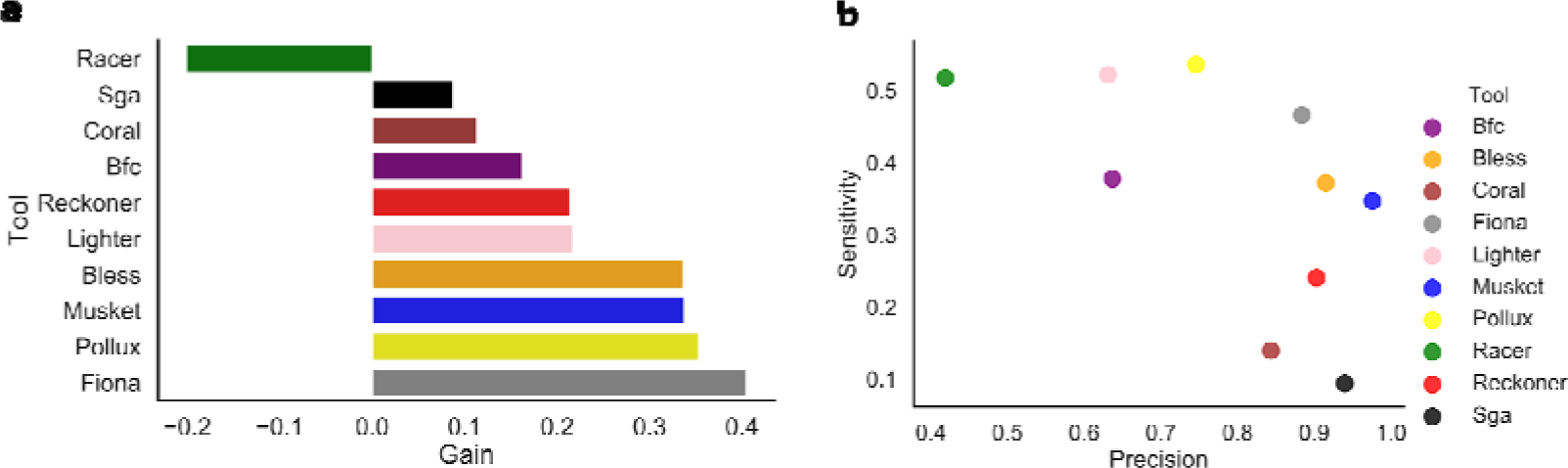
Correcting errors in viral sequencing data (D4 dataset). For all plots, the best k-mer size was selected. **(a)** Bar plot depicting the gain across various error correction methods. **(b)** Scatter plot depicting the sensitivity (x-axis) and precision (y-axis) of each tool.

We performed additional analysis to investigate the factors contributing to the performance of error correction tools on the viral sequencing data. We used a real HIV-1 sequencing benchmark^20^ composed of five HIV-1 subtype B haplotypes mixed *in-vitro* (**D5 dataset**) (**Table 1**). To prepare error-free reads we have applied haplotype-based error correction protocol able to eliminate sequencing errors by matching the read with the haplotype of origin. After the haplotype and reads were matched, the sequencing errors are corrected by replacing bases from reads with the based from the haplotype of origin. Details about the D5 dataset and haplotype-based error correction protocol are provided in the **Methods section**.

In contrast to the results generated from the D4 HIV dataset, the majority of error correction methods were unable to accurately correct errors (**Additional file 1: Fig. S19**). Notably, the gain was below 6% across all the methods. The vast majority of the error correction methods were able to maintain precision above 60%. However, none of the tools were able to achieve sensitivity above 20% (**Additional file 1: Fig. S20**).

We further investigated the factors which influenced the reduced performance on D5 HIV mixture dataset. First, we varied the diversity between haplotypes. We have generated three datasets each consisting of two haplotypes. The diversity was measured using the Hamming distance and varied between 5.94% and 0.02%. The reduced diversity between haplotypes had a positive effect for the majority of the error correction method, allowing seven out of 10 methods to achieve positive gain on the dataset with the lowest diversity (Hamming distance between haplotypes is 0.02%) (**Additional file 1: Fig. S21**).

We have also performed additional experiments to investigate the effect of the number of errors present in the data on the ability of methods to accurately correct errors. We have computationally changed the error rate of viral dataset D5 (**Methods section**). In total, we have obtained eight datasets with the error rate ranging from 10^−6^ to 3.3×10^−3^. In general, increased error rate had a negative impact on the ability of the majority of the methods to accurately correct errors. Tools, which maintained consistent performance across dataset with various error rates were Fiona and Racer. Notably, Racer was able to maintain gain above 70% across all datasets with various error-rates (**Additional file 1: Fig. S22**).

## Discussion

Our systematic assessment of currently available error correction tools highlights the advantages and limitations of computational error correction techniques across different types of datasets containing different levels of heterogeneity. We evaluated the performance of error correction algorithms on typical DNA sequencing data and highly heterogeneous data derived from human immune repertoires and intra-host viral populations. We observed large variability in the performance of error correction methods when applied to different types of datasets, with no single method performing best on all types of data. For example, the majority of surveyed methods deliver improved sequencing reads for datasets with coverage 8x or higher when applied to WGS human data. The variability in observed performance of error correction tools emphasizes the importance of benchmarking in order to inform the selection of an appropriate tool for any given data set and research question.

We observed that majority of the methods are capable of producing accurate results only for high coverage datasets, suggesting that depth of coverage is an important parameter when considering the choice of error correction tools. We determined that genomic coverage of 2x or higher was required for Coral to achieve substantially better reads in the WGS human data. Other tools require higher coverage to successfully correct sequencing errors. For example, seven out of ten tools are only able to successfully correct errors for coverage 4x or higher. A genomic coverage of 16x allows several methods to correct more than 90% of the errors with high precision. For example, Fiona was able to correct 98% of the errors with 94% precision. Our results suggest that genomic coverage for WGS human data should be taken into account when choosing the appropriate error correction tool. We also evaluated the effect of k-mer size on the accuracy of error correction tools. An increase in k-mer size typically offers an increase in accuracy of error correction when applied to both WGS and TCR-Seq data.

Our study found that performance of error correction methods vary substantially when applied to data across various domains of biology, with no single method performing best on all types of examined datasets. We noticed that error correction methods are useful in the field of immunogenomics, where multiple error correction methods may significantly improve results— even for extremely low coverage rates. These results suggest that computational error correction tools have potential to replace UMI-based error correction protocols. UMIs are commonly applied to data in immunogenomics studies in order to correct sequencing errors, but UMI-based error correction may have a negative impact on the coverage and this increases the cost per base of sequencing.

Similarly, error correction methods are useful to reduce the number of errors in heterogeneous viral populations. Three out of ten methods were able to significantly improve the viral sequencing reads with the gain exceeding 30%.

Our benchmarking study focused on benchmarking computational error correction tools. The evaluation of error correction on downstream analyses has been performed and published elsewhere^8^ and is beyond the scope of the current study. In future studies, we anticipate that additional knowledge about the structured properties of analyzed genomes will be used to develop bioinformatics tools that produce more accurate and reliable results. For example, structures of genomes from different organisms are shaped by epistasis resulting in co-dependence of different variants^30,31^. The incorporation of the effects of epistasis into error-correction methods may help researchers distinguish between real and artificial genomic heterogeneity and eventually result in a higher accuracy of error correction.

## Methods

### Running error correction tools

Error correction tools were run using the directions provided with each of the respective tools (**Additional file 1: Table S1**). Wrappers were then prepared in order to run each of the respective tools as well as create standardized log files. When running the tools, we chose the Illumina technology option and paired-end mode when possible. In cases where the paired-end option is not available (**Table 1**), we prepared single-end reads obtained from paired-end data by disregarding pairing information and treating each read from the pair as a single-end read. The computational pipeline to compare error correction methods is open source, free to use under the MIT license, and available at https://github.com/Mangul-Lab-USC/benchmarking.error.correction.

### Generating error-free reads using UMI-based clustering

Error-free reads for gold standard were generated using UMI-based clustering. Reads were grouped based on matching UMIs and corrected by consensus, where an 80% majority was required to correct sequencing errors without affecting naturally occurring SNVs (**Figure 1b**). UMI-based clustering was used to produce error-free reads for the D2 and D4 datasets.

### Generating simulated datasets

We generated simulated data mimicking the WGS data (D1 dataset) and TCR-Seq data (D3 dataset). To generate the D1 dataset, we developed a customized version of the tool WgSim^19^ **(Additional file 1: Fig. S1).** We simulated reads from chromosome 21. Read coverage varied between 1 and 32. Briefly, the customized version, along with generating the sequencing reads with errors, can report the error-free reads to the files provided as command line arguments. The WgSim fork is available at https://github.com/mandricigor/wgsim. Commands for generating the datasets are described in **Additional file 1: Supplemental Note 1.**

To generate the TCR-Seq dataset, we have used the T cell receptor alpha chain (TCRA)^33^. We generated samples with read lengths of 100bp. Read coverage varied between 1 and 32. For all the samples, the mean fragment length was set to 200 bp.

### Generating error-free reads using haplotype-based error correction protocol

We prepared viral dataset D5 using real sequencing data from NCBI with the accession number SRR961514 prepared by Giallonardo et al.^20^. This is a MiSeq sequencing experiment on a mixture of five subtype B HIV-1 viruses with different genomes. The original dataset contains 714994 MiSeq 2×250bp reads that we mapped on all five HIV-1 reference genomes. Each read was assigned to the reference with which it has a minimum number of mismatches. Since unmapped reads do not have the best match, we dropped them; as a result, there were 706182 remaining reads. The original error rate in the dataset was 1.44%. We modified these reads as follows: first, we corrected the corresponding portion of errors with a corresponding reference nucleotides to obtain different levels of errors in the datasets (1.44%, 0.33%, 0.1%, 0.033%, 0.01%, 0.0033%, 0.001%, 0.00033%, 0.0001%); second, we created datasets with mixtures of two haplotypes with the original 1.44% error rate but with different levels of diversity between haplotypes (Hamming distance=5.94%, 0.29%, 0.02%). Two haplotypes “89.6” and “YU2” were chosen from the original dataset SRR961514. The original haplotypes have the Hamming distance that equal 0.0595%. The random portion of “YU2” haplotype was corrected to reduce its distance to “89.6”. The MiSeq reads from “89.6” were corrected as well. We controlled that our correction did not fix sequencing errors. So, if the correction at a certain position of the read ended up in removing a sequencing error, we introduced it back at the same position by introducing random erroneous nucleotide.

### Error correction methods designed for mixed genomes

Most error correction methods are designed for a single genome, yet Pollux is a unique method designed for metagenomics data composed of multiple microbial genomes. It also can work for sequencing data derived from a single genome. Pollux determines the number of occurrences of each observed k-mer in the data. The k-mer counts are used to determine k-mer depth profile for each read and localize sequencing errors.

### Choosing k-mer size

We use k-mer sizes ranging from 20bp to 30bp for each of the datasets. In cases where the error correction tool was equipped with an option for the genome size, we provided the length of corresponding genome size. The genome size used for the T cell immune repertoire sequencing was 405,000 bp (total length of all simulated TCR transcripts present in the sample), while the whole genome sequencing size used was 46,709,983bp (length of chr21) for human, and 5,594,605bp (length of all chromosomes) for E.coli. The genome size used for the viral sequencing (HIV) was 9,181bp.

### Evaluating error correction accuracy

The evaluation of error correction involves obtaining the error-free reads, the original raw reads, and the original reads corrected by computational error correction tools. Reads are then compared using multiple sequence alignment. We used MUSCLE^34^ to perform multiple sequence alignment. Raw read represents the base before the error correction tool has been used. E.C. read represents the base after the error correction tool has been used. True read represents the correct base. True positive (TP) indicates a sequencing error was correctly changed. False negative (FN) indicates that either an error was ignored, or an error was incorrectly changed (**Additional file 1: Fig. S1**). False positive (FP) indicates a correct base was changed to an incorrect base. True negative (TN) indicates a correct base was left as is. Trimming was additionally evaluated as either TP or FP trimming. FN base calls were evaluated as either FN wrong if the base was changed incorrectly or just FN if the base was untouched and should have been corrected. (**Additional file 1: Fig. S2**). We have also reported CPU time and the maximum amount of RAM used by each of the tools.

### Data compression format

Due to the quantity and size of the error corrected fastq files, the evaluation of the reads was compressed. In order to summarize the errors that were not resolved by each of the various tools, a method similar to the evaluation of error correction was utilized. In substitute for determining the number of TP, TN, FP from INDELs, FP from trimming, normal FP, and FN bases, the data compression will represent this data in the following reduced manner. The format is in the following order: read_name, length, TP, FN, FN WRONG, FP, FP INDEL, FP TRIM, TP TRIM (Example: 1_22_238_1:0:0_3:0:0_0/1,100,3,0,0,0,0,0,0). By producing this data only when a TP, FP of any type, and FN are encountered.

### Estimating performance

We compared the performance of the error correction tools by reporting wall time, CPU time, and the maximum amount of RAM used by each tool. These performance metrics were obtained via -qsub option, with an additional -m bse option allowing automatically generated CPU and memory usage statistics. A typical node of the cluster used to benchmark the tools has dual twelve-core 2.2GHz Intel ES-2650v4 CPUs and an Intel 800GB DC S3510 Series MLC (6 Gb/s, 0.3 DWPD) 2.5” SATA SSD.

### Comparing performance of tools across the genomic categories

We compared the performance of error correction tools across different genomic categories based on sequence complexity. In order to annotate genome (more precisely, chromosome 21 of the human genome) with a category, we used RepeatMasker (version 4.0.9). As a result, the genome was divided into multiple categories (the most abundant ones are: “LINE/L1”, “SINE/Alu”, “LTR/ERVL-MaLR”, “LINE/L2”, “LTR/ERV1”, “LTR/ERVL”, “SINE/MIR”, “Simple_repeat”, “DNA/hAT-Charlie”, “DNA/TcMar-Tigger”, “Satellite/centr”, “DNA/hAT-Tip100”, “LTR/Gypsy”, “Low_complexity”, “LINE/CR1”, “LINE/RTE-X”, “Satellite”, “LTR”, “LTR/ERVK”). We also introduced a category “normal” which consists of sequences not in any of the aforementioned categories. A read is considered to belong to a category X if it overlaps a sequence from category X.

## Authors’ contributions

K.M. created scripts for running and evaluating software tools, developed methods and contributed to writing the manuscript. J.J.B. contributed to creating scripts for running and evaluating software tools on a cluster, generating figures, and writing of the manuscript. I.M. developed methods and evaluated software tools. Q.W., S.C., R.L., B.L.H, K.H., E.L., G.E., D.Y., and P.S. contributed to installing, running and evaluating software tools. S.K., N.C.W., and R.S. contributed to the generation of datasets. L.S.M. and A.K. contributed to the production of figures and writing of the manuscript. E.G., H.Y., L.C., T.S., J.S., and M.P contributed to writing the manuscript. E.E. and A.Z. contributed to writing and method design. S.M. led the project and contributed to writing the manuscript.

## Acknowledgments

We thank the UCLA Bruins-In-Genomics (B.I.G.) Summer Research Program for supporting undergraduate researchers who contributed to this paper.

## Ethics approval and consent to participate

Ethics approval was not applicable

## Availability of data and materials

The D1 and D3 datasets were produced by computational simulations using a customized version of the WgSim tool available at https://github.com/mandricigor/wgsim. SRA data was downloaded via SRA archive (https://www.ncbi.nlm.nih.gov/sra). The 8 samples of the D2 dataset correspond to the accession numbers: SRR1543964, SRR1543965, SRR1543966, SRR1543967, SRR1543968, SRR1543969, SRR1543970, and SRR1543971. D4 dataset is identified by the accession number SRR11207257, which corresponds to a UMI-based HIV population sequencing of an infected patient, available at https://www.ncbi.nlm.nih.gov/sra/SRR11207257. D5 dataset was generated using real sequencing data from NCBI with the accession number SRR961514. Raw and true reads for datasets D1, D2, D3, D4, and D5 are available at https://dx.doi.org/10.6084/m9.figshare.11776413. The computational pipeline to compare error correction methods is open source, and publicly available at https://github.com/Mangul-Lab-USC/benchmarking.error.correction^32^ under the MIT license.

## Additional files

### Additional file 1.pdf

**Figure S1.** Flowchart displaying the modification made to WgSim. **Figure S2.** Various scenarios of error correction at the base level. **Figure S3.** Methodology to evaluate the accuracy of error correction methods. **Figure S4.** The effect of the k-mer size on the accuracy of the total corrections (D1 dataset). **Figure S5.** The effect of coverage on the total number of corrections (D1 dataset). **Figure S6.** The effect of coverage on the accuracy of the tools (D1 dataset). **Figure S7.** The portion of trimmed bases across various coverages settings (D1 dataset). **Figure S8.** The efficiency of trimming across various coverages settings (D1 dataset). **Figure S9.** Trimming Efficiency vs. Trim Portion (D1 dataset). **Figure S10.** Heatmap depicting the gain for low and normal complexity regions (D1 dataset). **Figure S11.** Barplot depicting the CPU time and the maximum amount of RAM across tools (D1 dataset). **Figure S12**. The effect of k-mer size on the accuracy of the tools (D3 dataset). **Figure S13.** The effect of k-mer size on the accuracy of the tools (D2 dataset). **Figure S14.** Heatmap depicting the gain across various coverage settings of TCR-seq (D3 dataset). **Figure S15.** Heatmap depicting the precision across various coverage settings of TCR-seq (D3 dataset). **Figure S16.** Heatmap depicting the sensitivity across various coverage settings of TCR-seq (D3 dataset). **Figure S17.** Trimming Efficiency vs. Trim Portion for TCR-Seq simulated data (D3 dataset). **Figure S18.** The effect of k-mer size on the accuracy of the tools (D4 dataset). **Figure S19.** Barplot depicting the gain across various error correction methods when applied to D5 HIV mixture dataset. **Figure S20.** Scatter plot depicting the sensitivity and precision of each tool for D5 HIV mixture dataset **Figure S21.** Heatmap depicting the gain across D5 HIV mixture dataset for various rates of diversity between haplotypes. **Figure S22.** Heatmap depicting the gain across D5 HIV mixture dataset with various error rates. **Table S1.** Instructions for running the tools. **Table S2.** Best k-mer size for each tool for D1 dataset. **Table S3.** Best k-mer size for each tool for D2 dataset. **Table S4.** Best k-mer size for each tool for D4 dataset. **Table S5.** Best k-mer size for each tool for D4 dataset. **Table S6.** Best k-mer size for each tool for D5 dataset.

The review history is available as **Additional file 2.pdf**.

## References Cited

1. Schuster, S. C. Next-generation sequencing transforms today’s biology. Nature Methods vol. 5 16–18 (2008).

2. Scholz, M. B., Lo, C.-C. & Chain, P. S. G. Next generation sequencing and bioinformatic bottlenecks: the current state of metagenomic data analysis. Curr. Opin. Biotechnol. 23, 9–15 (2012).

3. Salk, J. J., Schmitt, M. W. & Loeb, L. A. Enhancing the accuracy of next-generation sequencing for detecting rare and subclonal mutations. Nat. Rev. Genet. 19, 269–285 (2018).

4. Ma, X. et al. Analysis of error profiles in deep next-generation sequencing data. Genome Biol. 20, 50 (2019).

5. Strom, S. P. Current practices and guidelines for clinical next-generation sequencing oncology testing. Cancer Biol Med 13, 3–11 (2016).

6. Robasky, K., Lewis, N. E. & Church, G. M. The role of replicates for error mitigation in next-generation sequencing. Nat. Rev. Genet. 15, 56–62 (2014).

7. Ratan, A. et al. Comparison of Sequencing Platforms for Single Nucleotide Variant Calls in a Human Sample. PLoS One 8, e55089 (2013).

8. Heydari, M., Miclotte, G., Demeester, P., Van de Peer, Y. & Fostier, J. Evaluation of the impact of Illumina error correction tools on de novo genome assembly. BMC Bioinformatics 18, 374 (2017).

9. Liu, Y., Schröder, J. & Schmidt, B. Musket: a multistage k-mer spectrum-based error corrector for Illumina sequence data. Bioinformatics 29, 308–315 (2013).

10. Heo, Y., Wu, X.-L., Chen, D., Ma, J. & Hwu, W.-M. BLESS: bloom filter-based error correction solution for high-throughput sequencing reads. Bioinformatics 30, 1354–1362 (2014).

11. Marinier, E., Brown, D. G. & McConkey, B. J. Pollux: platform independent error correction of single and mixed genomes. BMC Bioinformatics 16, 10 (2015).

12. Chen, Z. et al. Highly accurate fluorogenic DNA sequencing with information theory-based error correction. Nat. Biotechnol. 35, 1170–1178 (2017).

13. Yang, X., Chockalingam, S. P. & Aluru, S. A survey of error-correction methods for next-generation sequencing. Brief. Bioinform. 14, 56–66 (2013).

14. Molnar, M. & Ilie, L. Correcting Illumina data. Brief. Bioinform. 16, 588–599 (2015).

15. Mangul, S. et al. Systematic benchmarking of omics computational tools. Nat. Commun. 10, 1393 (2019).

16. Laehnemann, D., Borkhardt, A. & McHardy, A. C. Denoising DNA deep sequencing data— high-throughput sequencing errors and their correction. Brief. Bioinform. 17, 154–179 (2015).

17. Zhang, T.-H., Wu, N. C. & Sun, R. A benchmark study on error-correction by read-pairing and tag-clustering in amplicon-based deep sequencing. BMC Genomics 17, 108 (2016).

18. Kinde, I., Wu, J., Papadopoulos, N., Kinzler, K. W. & Vogelstein, B. Detection and quantification of rare mutations with massively parallel sequencing. Proc. Natl. Acad. Sci. U. S. A. 108, 9530–9535 (2011).

19. lh. lh3/wgsim. GitHub https://github.com/lh3/wgsim.

20. Giallonardo, F. D. et al. Full-length haplotype reconstruction to infer the structure of heterogeneous virus populations. Nucleic Acids Res. 42, e115 (2014).

21. Salmela, L. & Schröder, J. Correcting errors in short reads by multiple alignments. Bioinformatics 27, 1455–1461 (2011).

22. Schulz, M. H. et al. Fiona: a parallel and automatic strategy for read error correction. Bioinformatics 30, i356–63 (2014).

23. Li, H. BFC: correcting Illumina sequencing errors. Bioinformatics 31, 2885–2887 (2015).

24. Song, L., Florea, L. & Langmead, B. Lighter: fast and memory-efficient sequencing error correction without counting. Genome Biol. 15, 509 (2014).

25. Ilie, L. & Molnar, M. RACER: Rapid and accurate correction of errors in reads. Bioinformatics 29, 2490–2493 (2013).

26. Dlugosz, M. & Deorowicz, S. RECKONER: read error corrector based on KMC. Bioinformatics 33, 1086–1089 (2017).

27. Simpson, J. T. & Durbin, R. Efficient de novo assembly of large genomes using compressed data structures. Genome Res. 22, 549–556 (2012).

28. Wirawan, A., Harris, R. S., Liu, Y., Schmidt, B. & Schröder, J. HECTOR: a parallel multistage homopolymer spectrum based error corrector for 454 sequencing data. BMC Bioinformatics vol. 15 (2014).

29. Olson, D. L. & Delen, D. Advanced Data Mining Techniques. (Springer Science & Business Media, 2008).

30. Diament, A. & Tuller, T. Tracking the evolution of 3D gene organization demonstrates its connection to phenotypic divergence. Nucleic Acids Res. 45, 4330–4343 (2017).

31. Shi, Y. et al. Chromatin accessibility contributes to simultaneous mutations of cancer genes. Sci. Rep. 6, 35270 (2016).

32. Mitchell, K. et al. Repository for our benchmarking study ‘Benchmarking of computational error-correction methods for next-generation sequencing’. GitHub https://github.com/Mangul-Lab-USC/benchmarking_error_correction (2019).

33. Mangul, S. et al. Profiling immunoglobulin repertoires across multiple human tissues by RNA Sequencing. doi: 10.1101/089235.

34. Edgar, R. C. MUSCLE: multiple sequence alignment with high accuracy and high throughput. Nucleic Acids Res. 32, 1792–1797 (2004).

