## Supplementary Materials for "Benchmarking of computational error-correction methods for next-generation sequencing data"

**Additional file 1**

**Supplemental Note 1: Commands to generate simulated datasets**

We simulated WGS data from chromosome 21 using the following command:

wgsim -r 0.001 -R 0.0001 -e 0.005 -1 $rlen -2 $rlen -A 0 -N $nr Escherichia_coli_str_k_12_substr_mg1655.ASM584v2.dna.chromosome.Chromosome.fa datasets/ecoli/wgsim_rl_${rlen}_cov_${cov}.1.fastq datasets/ecoli/wgsim_rl_${rlen}_cov_${cov}.2.fastq true_reads/ecoli/true_rl_${rlen}_cov_${cov}.1.fastq true_reads/ecoli/true_rl_${rlen}_cov_${cov}.2.fastq 1>logfile.txt 2>err.txt

**Supplemental Note 2: Preparing data for evaluation**

We have used the first 200 thousand reads to evaluate the performance of the error correction methods on HIV mixture data (D5 dataset) with varying error rates **(Fig. S22).** For running Bless, we have removed reads smaller than 30bp from all datasets as it fails to run on reads smaller than the k-mer size specified.

### **Supplementary Figures**

Generating error-free reads

Generating alternative reference

Introducing errors into the reads

Print out reads with errors to a file

Print out error-free reads to a file

**Flowchart of our customized version of wgsim tool.**

The red box stands for the updates we introduced into wgsim to print out error free reads.

**Figure S1.** Flowchart displaying the modification made to WgSim in order to produce error-free reads (highlighted in red) in addition to the reads with the errors.

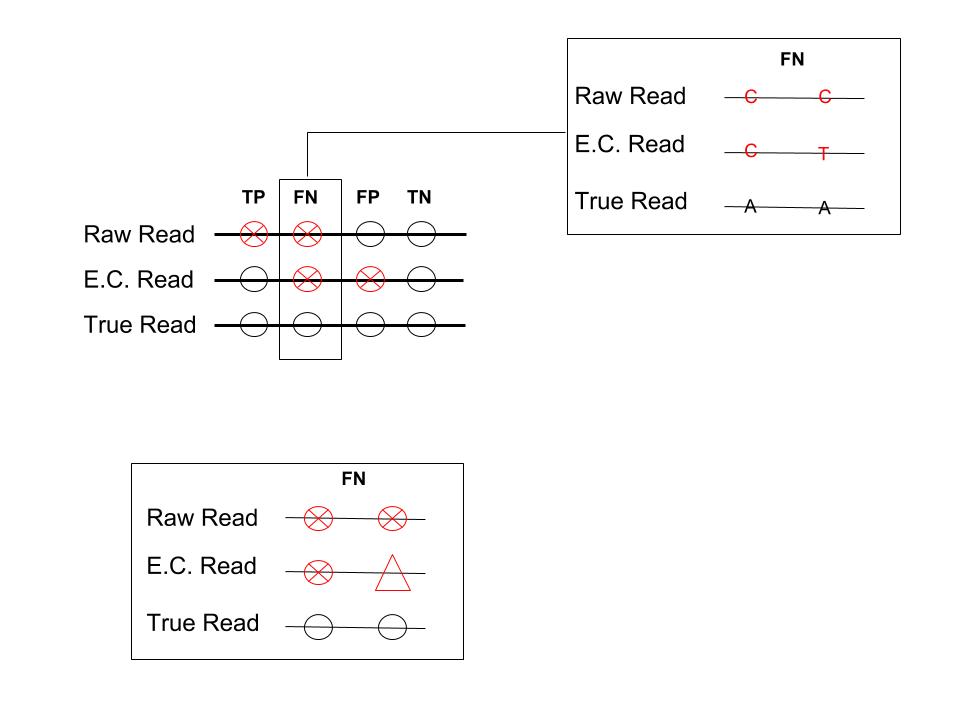

**Figure S2.** Various scenarios of error correction at the base level. Raw read represents the base before the error correction tool has been used. E.C. read represents the base after the error correction tool has been used. True read represents the correct base. True positive (TP) indicates a sequencing error was correctly changed. False negative (FN) indicates that either an error was ignored, or an error was incorrectly changed. False positive (FP) indicates a correct base was changed to an incorrect base. True negative (TN) indicates a correct base was left as is.

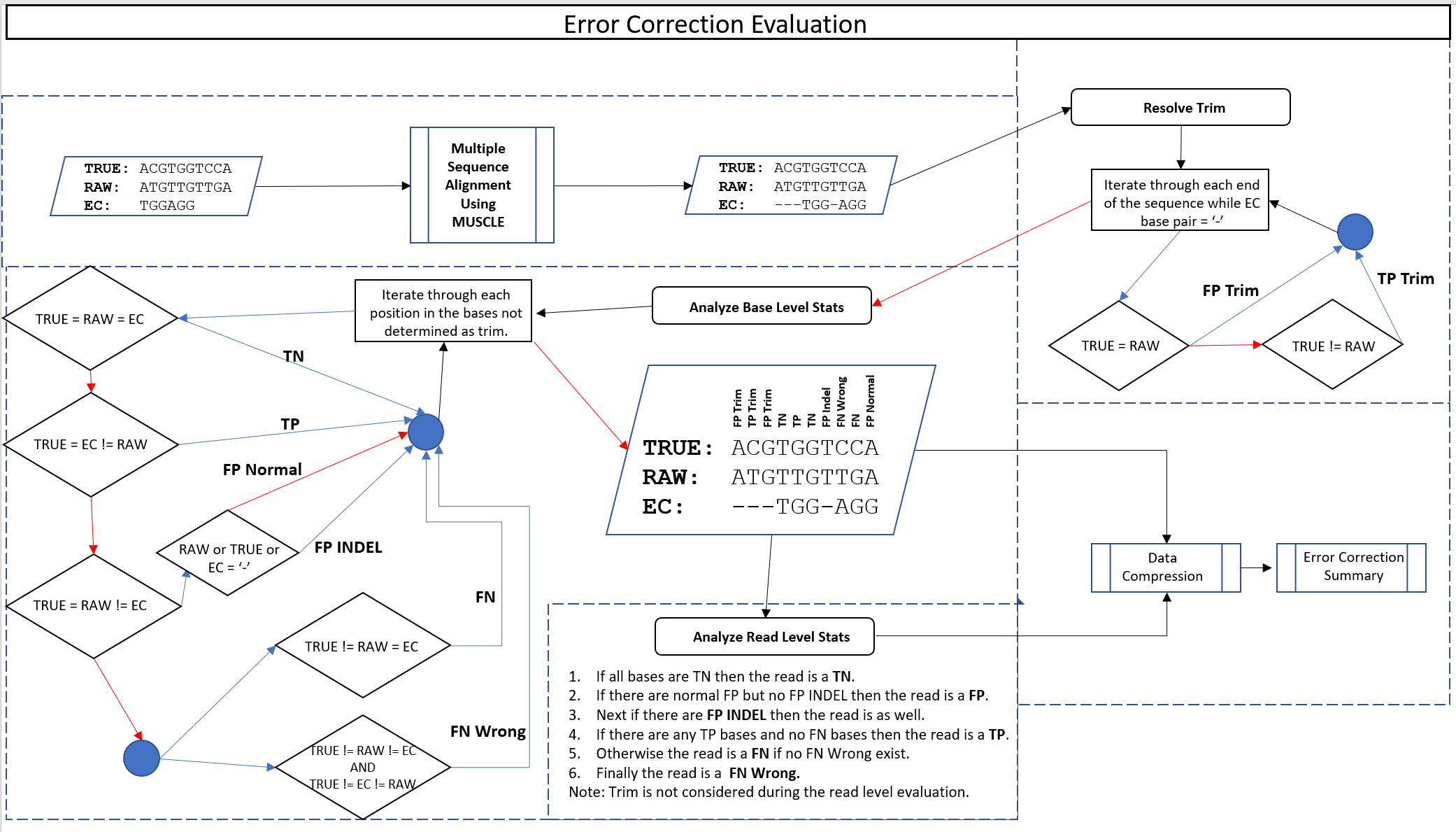

**Figure S3.** Methodology to evaluate the accuracy of error correction methods. The image above displays the series of events that take place in order to progress from the error-corrected read supplied from any one of the tools mentioned in Table 2***.*** MSA is first performed to enable an analysis of each base in comparison to the “gold standard”, or proposed true read, as well as the raw read prior to error correction. The progression of events following MSA is Resolve Trim, Analyze Base Level Stats, Analyze Read Level Stats, Data Compression, and finally the Error Correction Summary. Algorithmic details for Resolving Trim, Analyze Base Level Stats, and Analyze Read Level Stats are shown above within each of the relevant dashed boxes. Red or blue arrows represent whether a decision was false or true; in addition, a red or blue arrow represents whether a process iteration is over or still in progress.

**(a)**

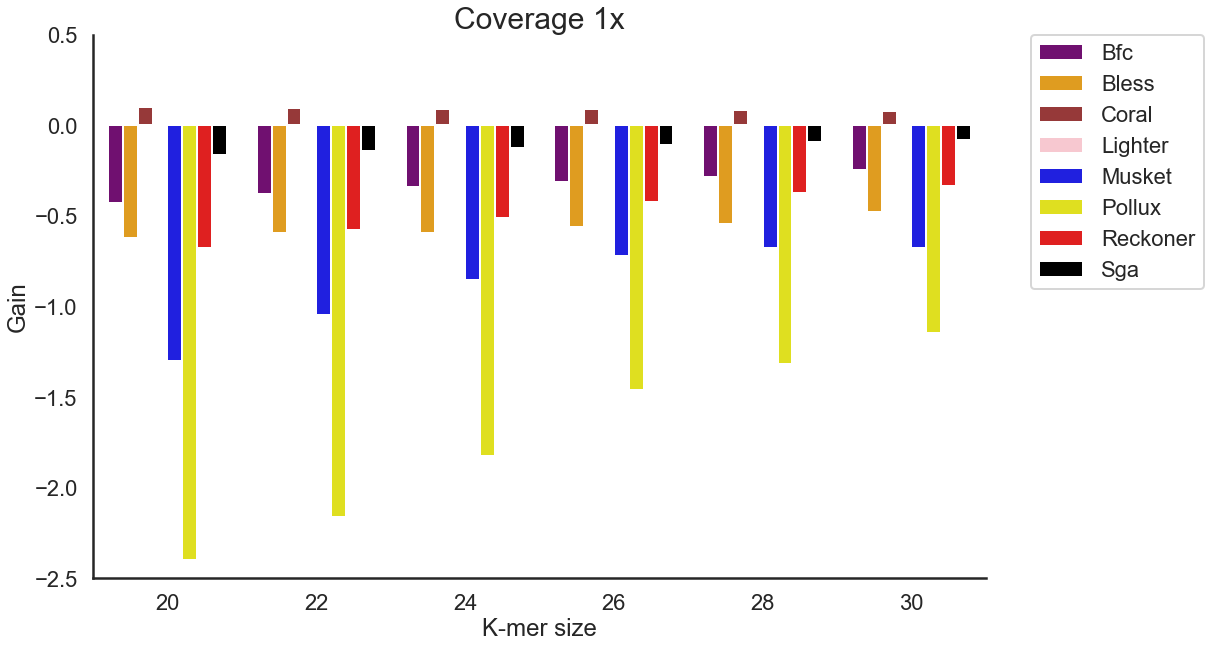

**(b)**

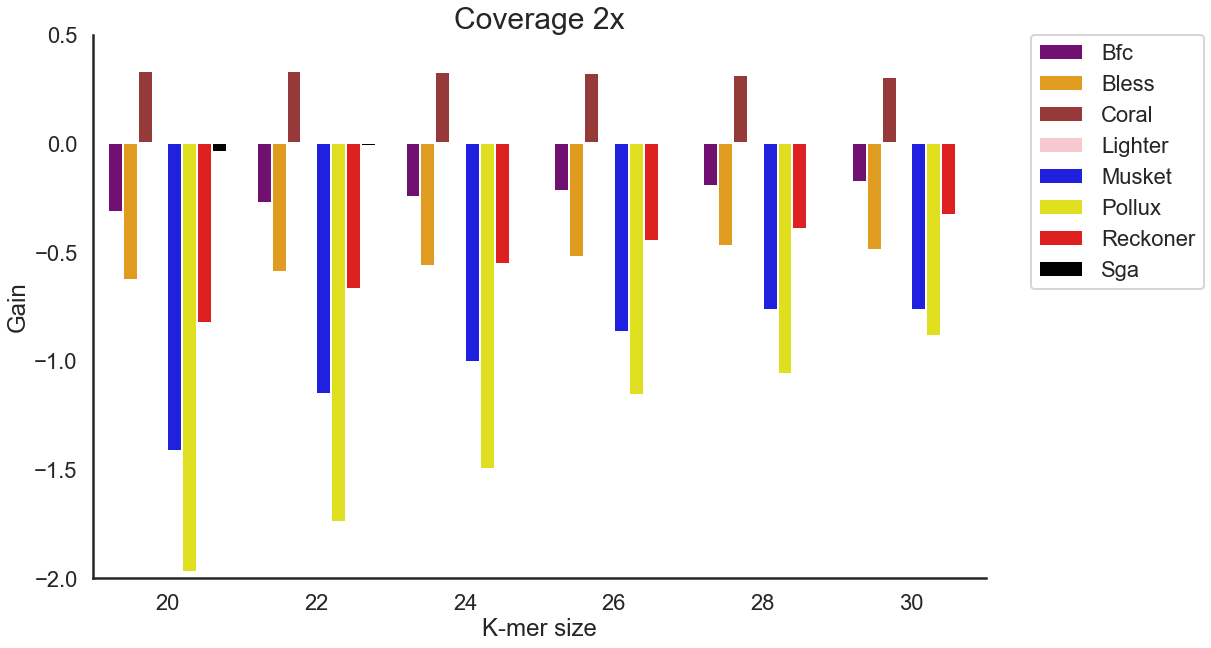

**(c)**

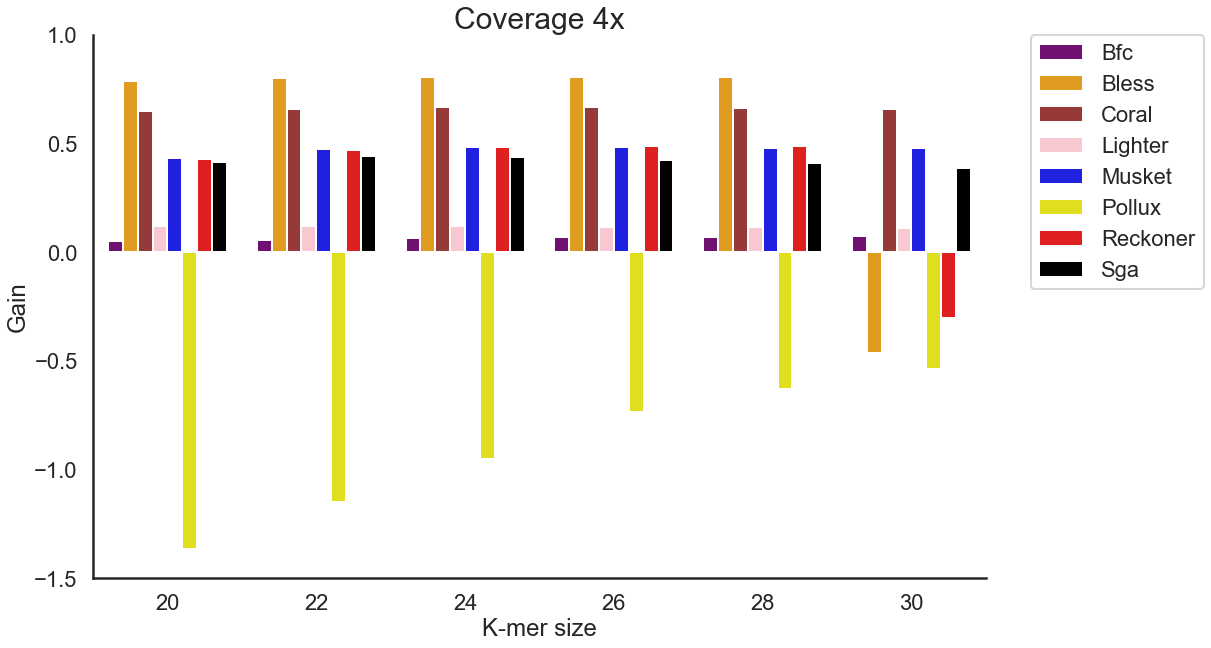

**(d)**

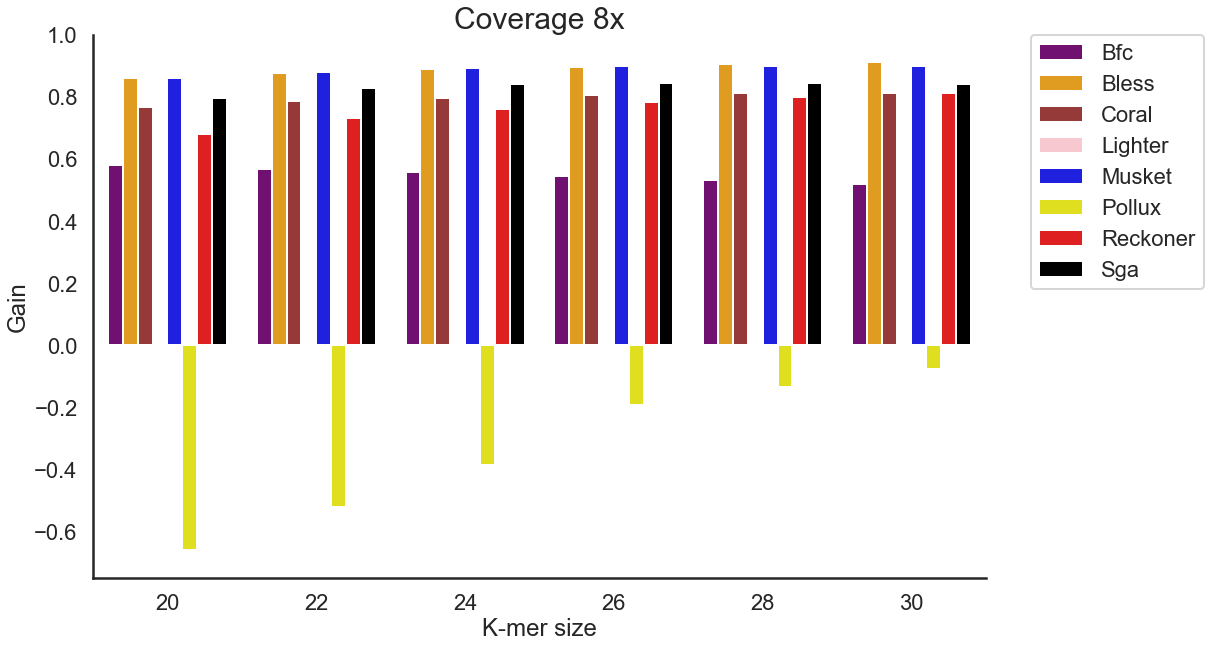

**(e)**

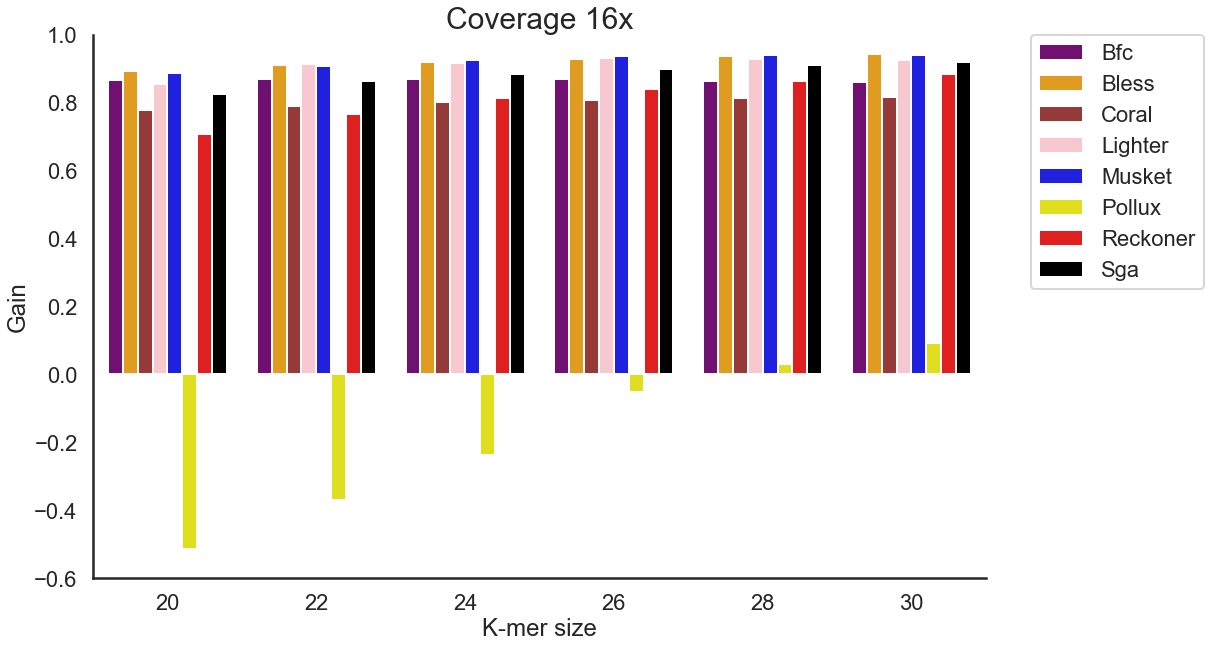

**(f)**

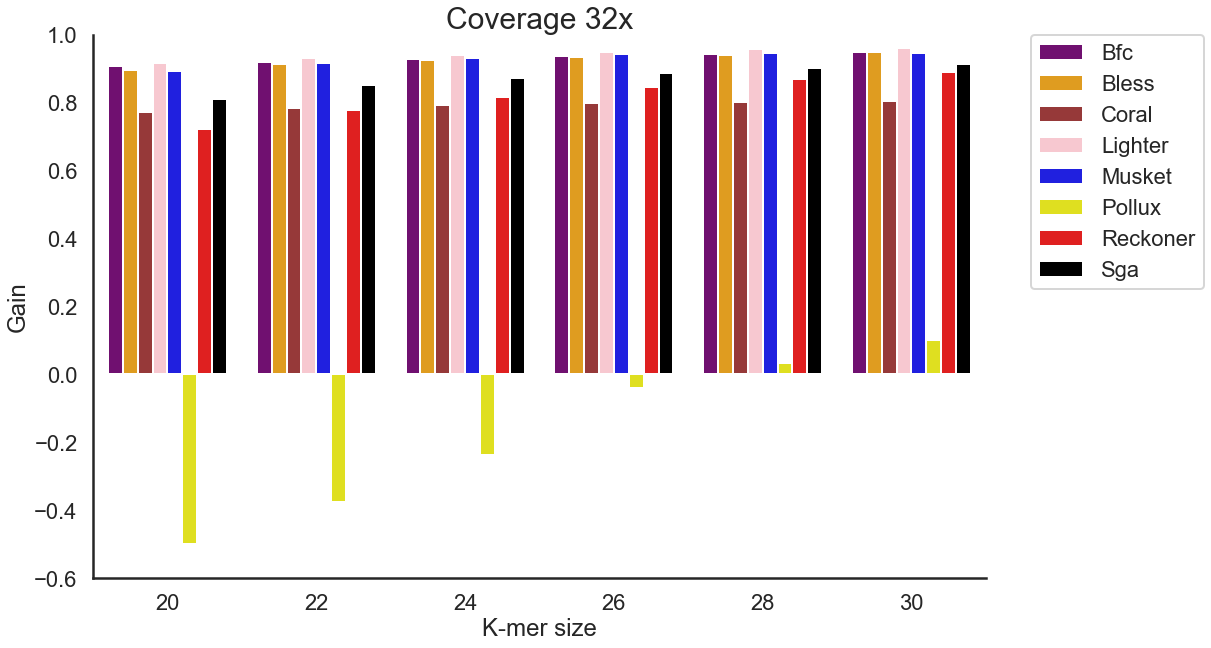

**Figure S4.** The effect of the k-mer size on the accuracy of the total corrections made across various coverages settings for WGS human data (D1dataset). (a) Coverage of 1x. (b) Coverage of 2x. (c) Coverage of 4x. (d) Coverage of 8x. (e) Coverage of 16x. (f) Coverage of 32x. We have excluded Fiona and Racer, as those tools do not provide options for k-mer size.

(a)
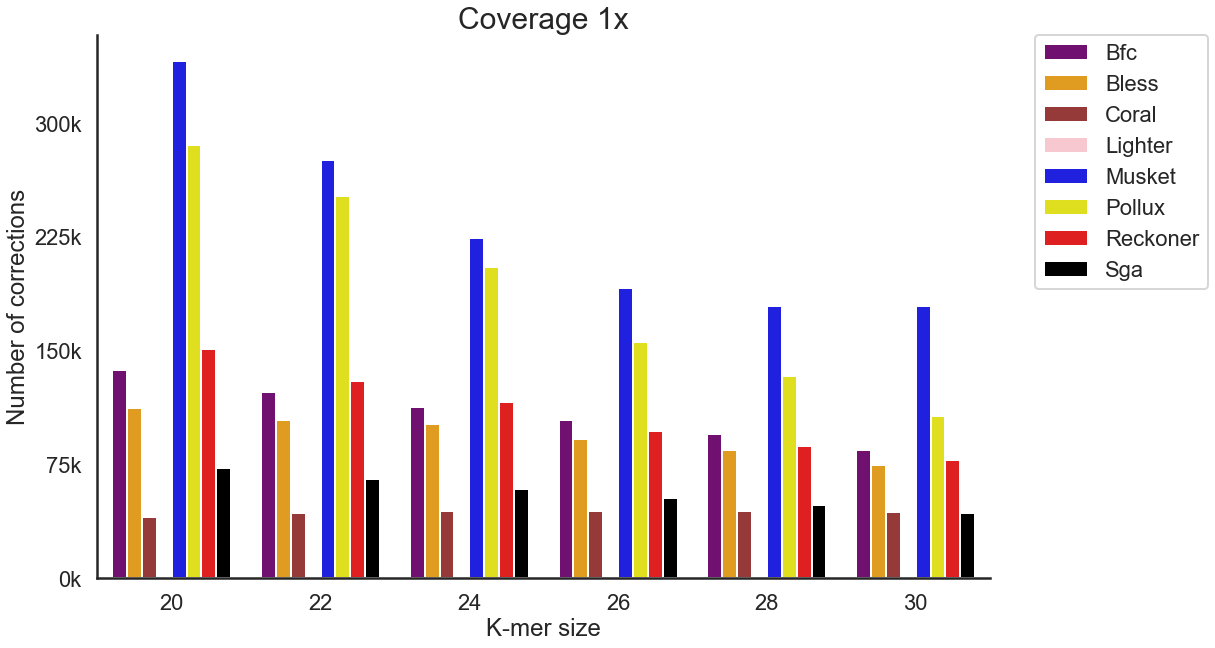

(b)

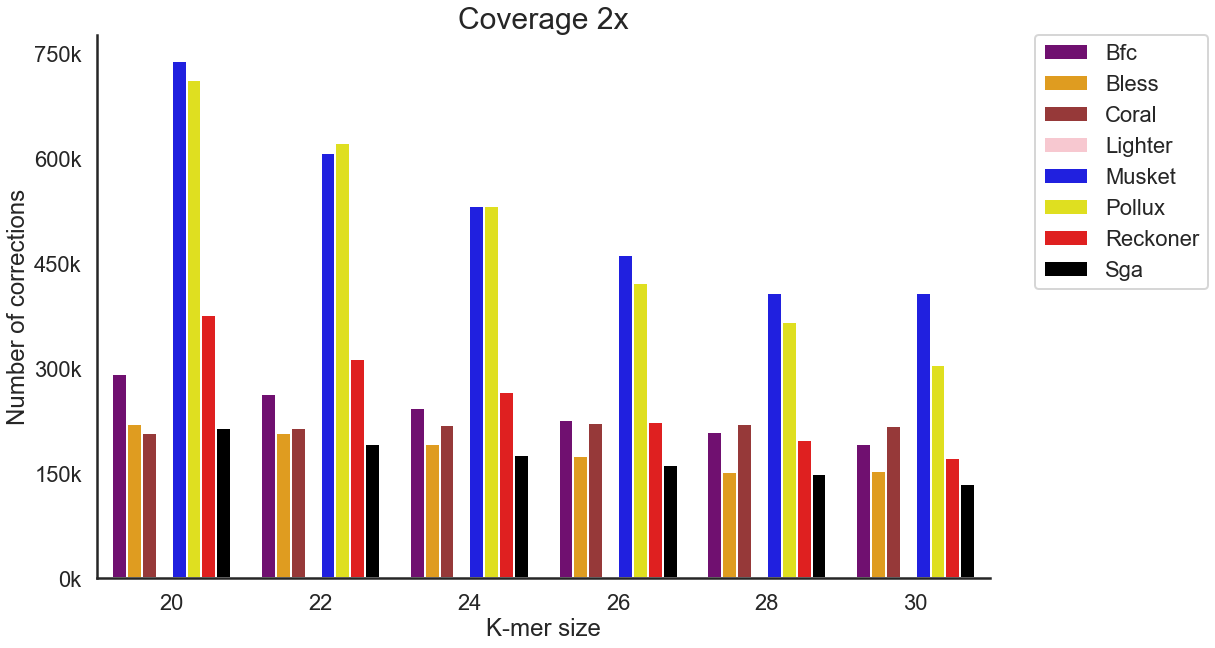

(c)

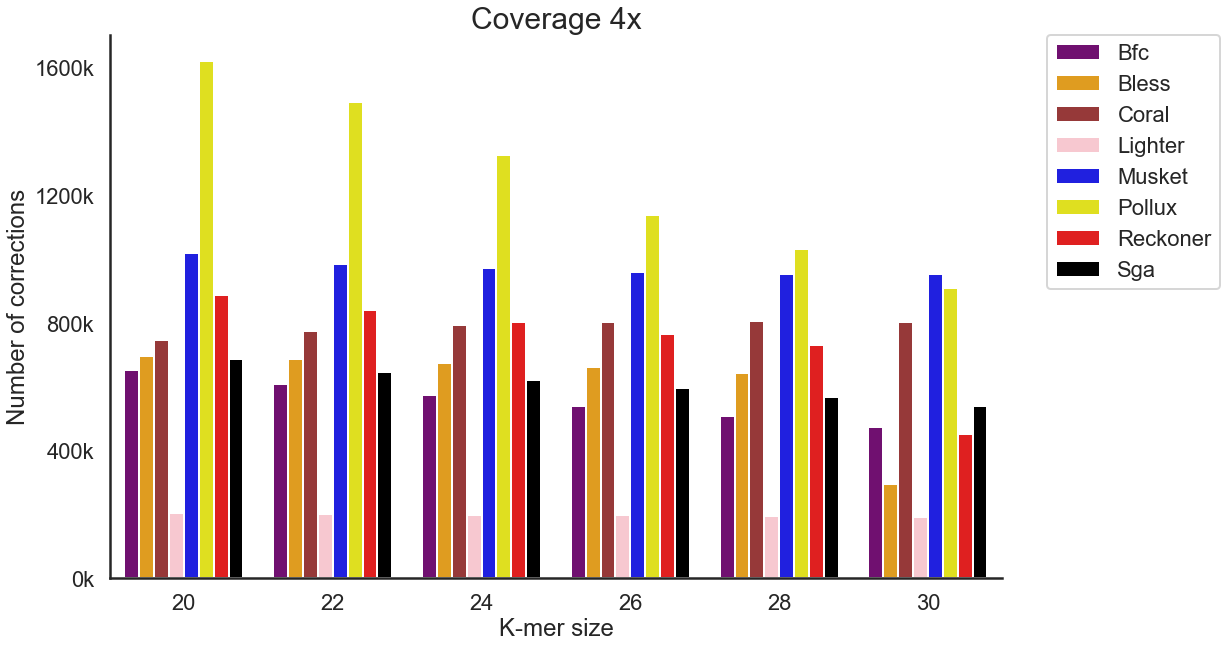

(d)

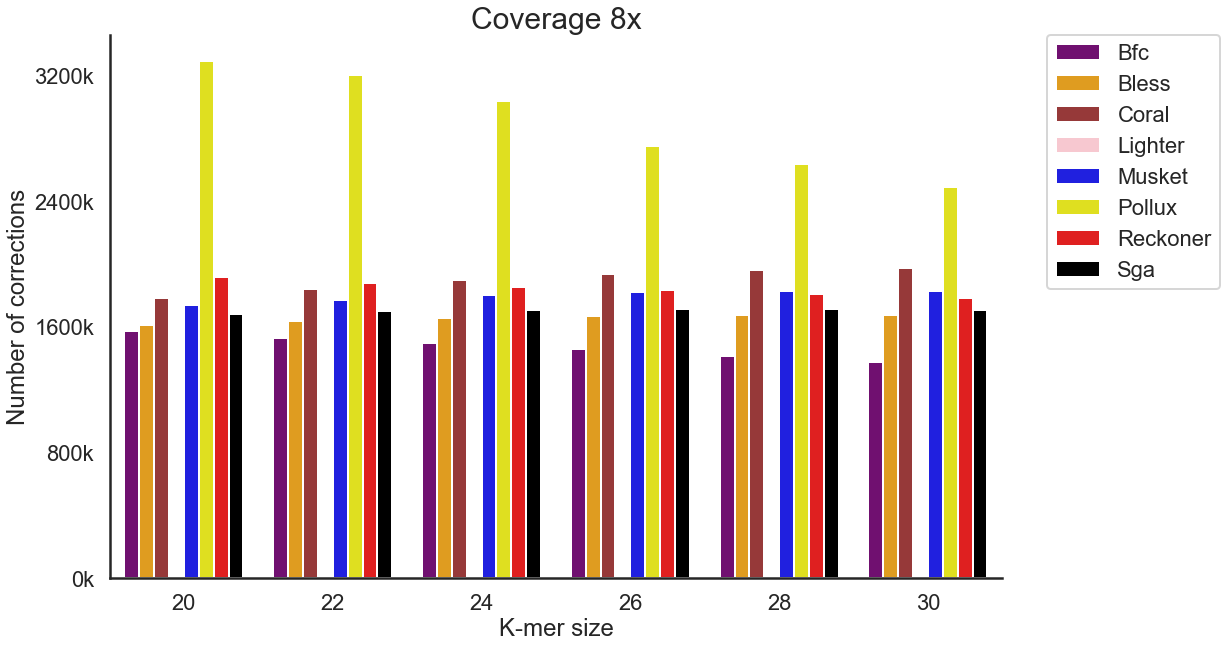

(e)

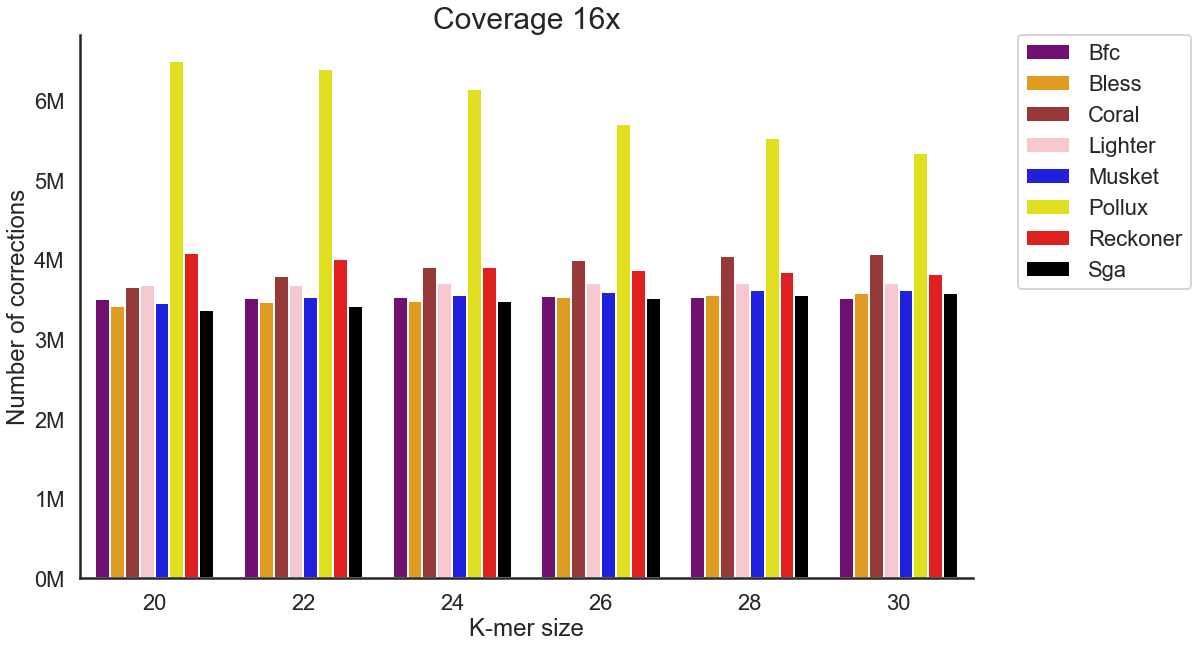

(f)

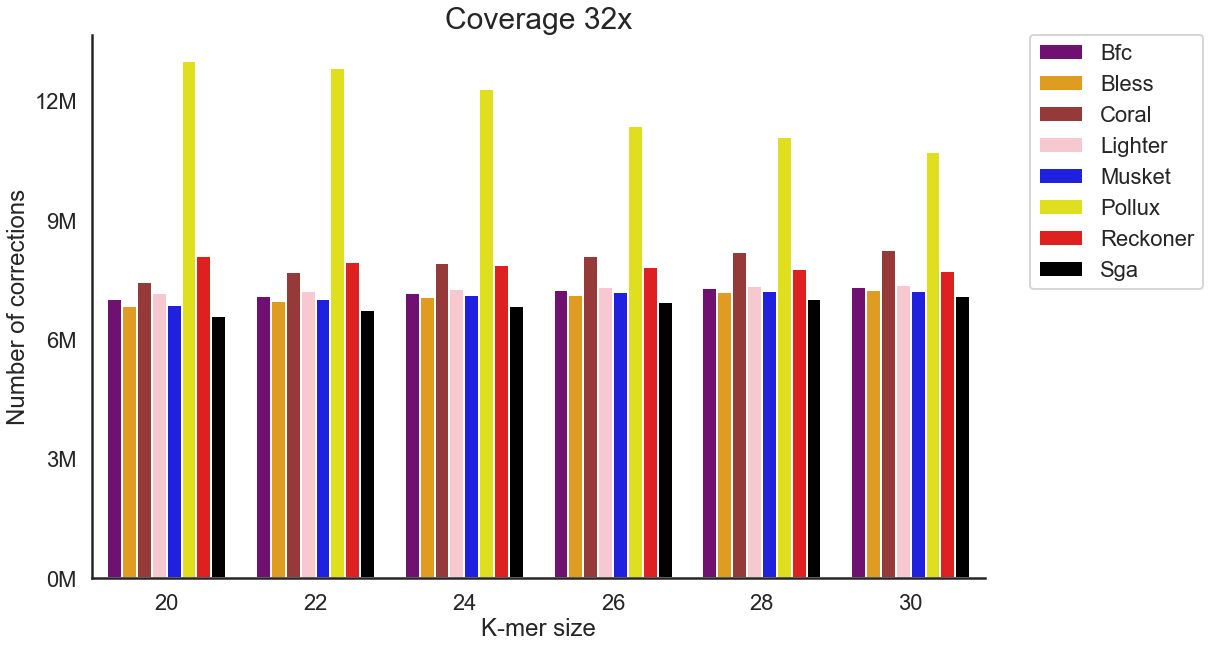

**Figure S5.** The effect of coverage on the total number of corrections performed by error correction methods across WGS human data (D1 dataset). (a) Coverage of 1x. (b) Coverage of 2x. (c) Coverage of 4x. (d) Coverage of 8x. (e) Coverage of 16x. (f) Coverage of 32x. We have excluded Fiona and Racer, as those tools do not provide options for k-mer size.

(a)

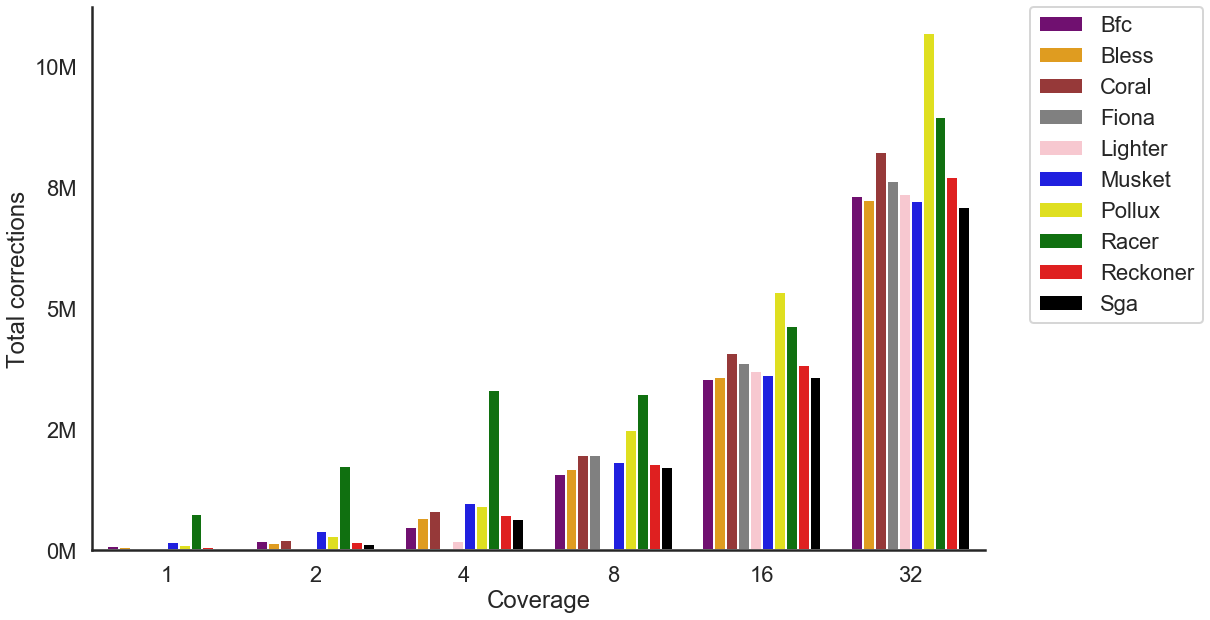

(b)

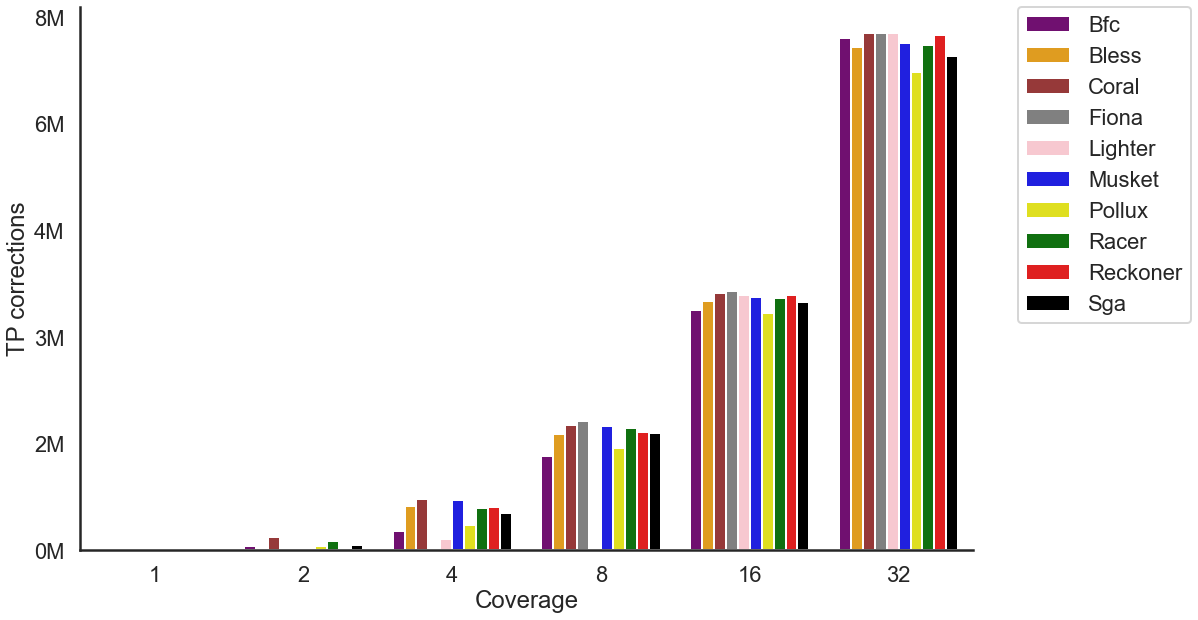

(c)

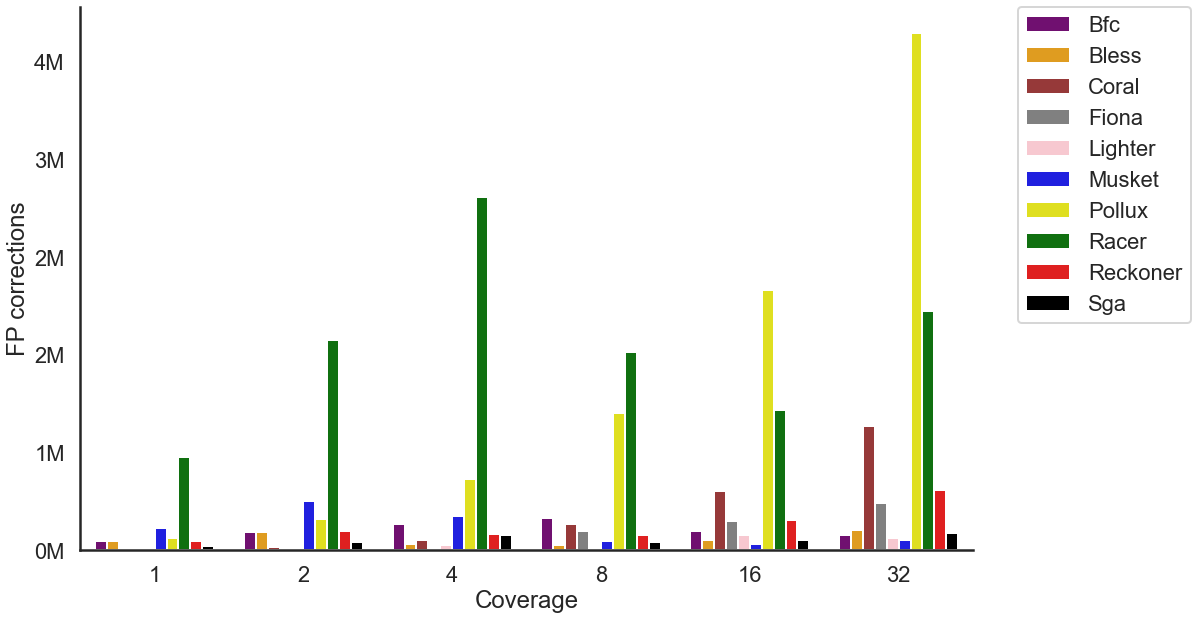

**Figure S6.** The effect of coverage on the accuracy of the error correction tools across various coverages settings for WGS human data (D1 dataset). (a) Total number of corrections. (b) Number of TP corrections. (c) Number of FP corrections. For each tool, the best k-mer size was selected.

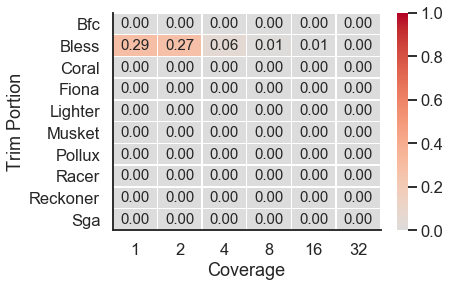

**Figure S7.** The portion of trimmed bases across various coverages settings for WGS human data (D1 dataset). For each tool, the best k-mer size was selected.

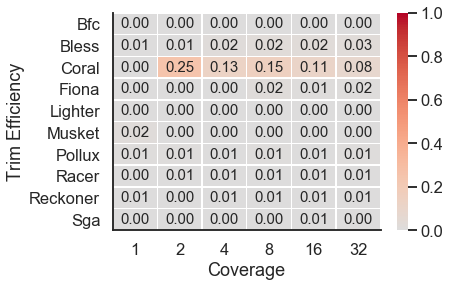

**Figure S8.** The efficiency of trimming across various coverages settings for WGS human data (D1 dataset). For each tool, the best k-mer size was selected.

(a)

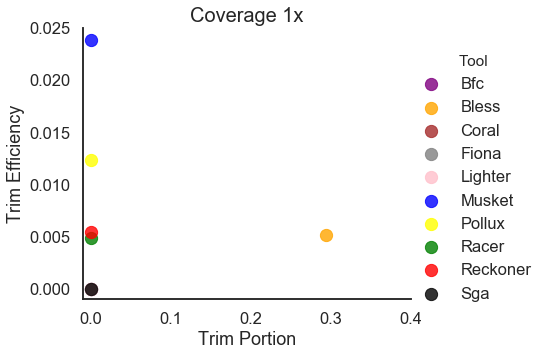

(b)

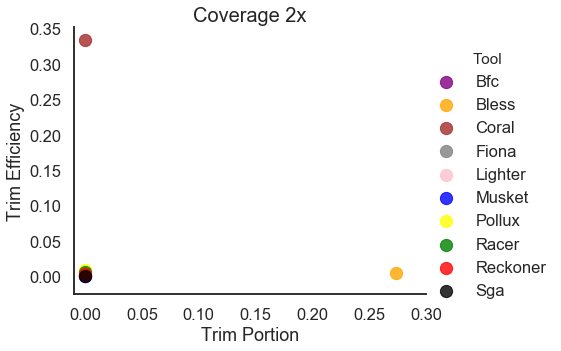

(c)

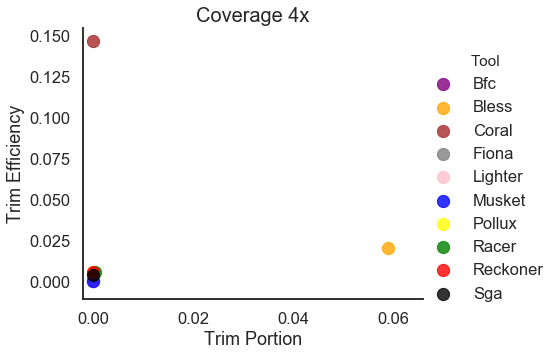

(d)

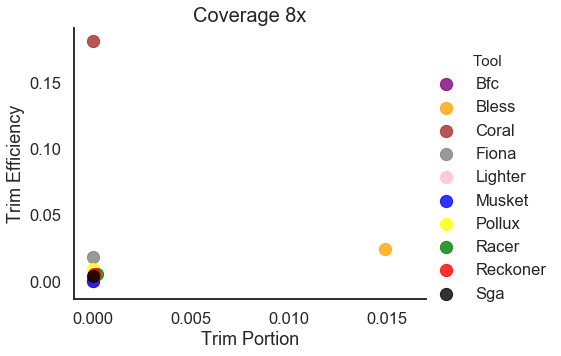

(e)

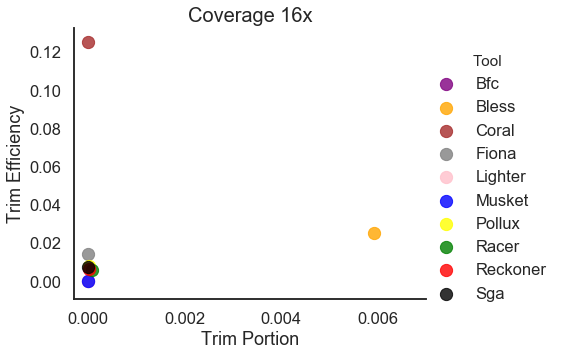

(f)

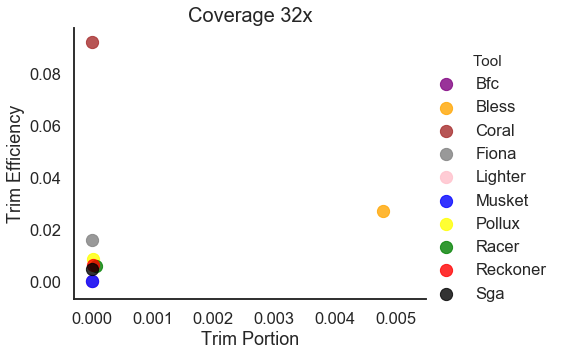

**Figure S9.** Trimming Efficiency vs. Trim Portion for WGS human data (D1 dataset). (a) Coverage of 1x. (b) Coverage of 2x. (c) Coverage of 4x. (d) Coverage of 8x. (e) Coverage of 16x. (f) Coverage of 32x. For each tool, the best k-mer size was selected.

(a)

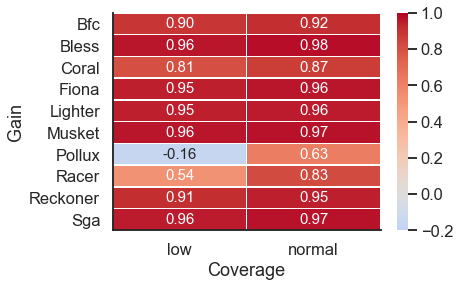

(b)

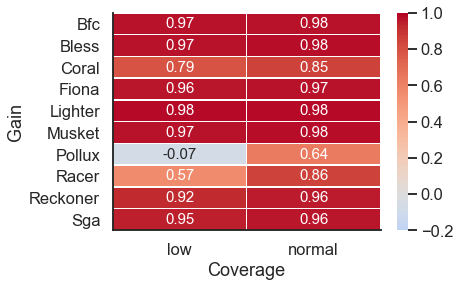

##### **Figure S10.** Heatmap depicting the gain for low complexity regions (‘low’) and the rest of the genome (‘normal’). (a) WGS human data (D1 dataset) with 16x coverage. (b) WGS human data with 32x coverage. For each tool, the best k-mer size was selected.

(a)

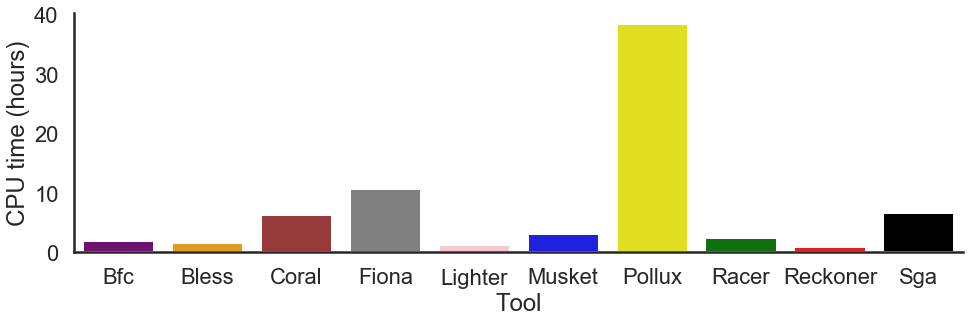

(b)

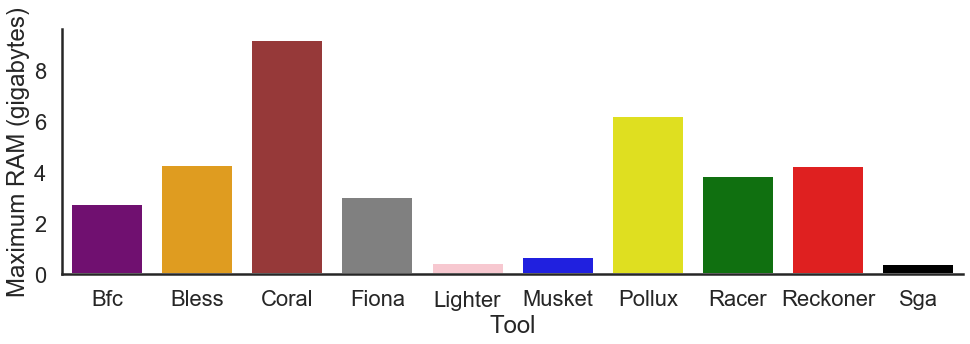

**Figure S11.** Barplot depicting (a) the CPU time and (b) the maximum amount of RAM across error correction tools for WGS human data (D1 dataset). For each tool, the mean value across different k-mer sizes was selected.

(a)

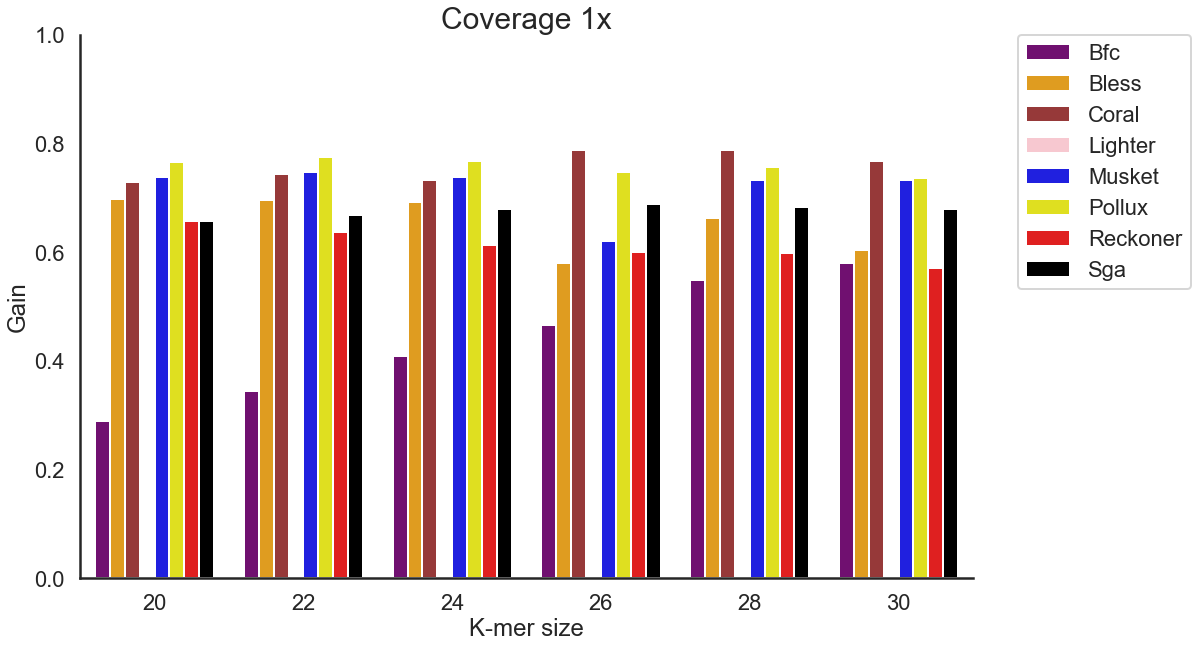

(b)

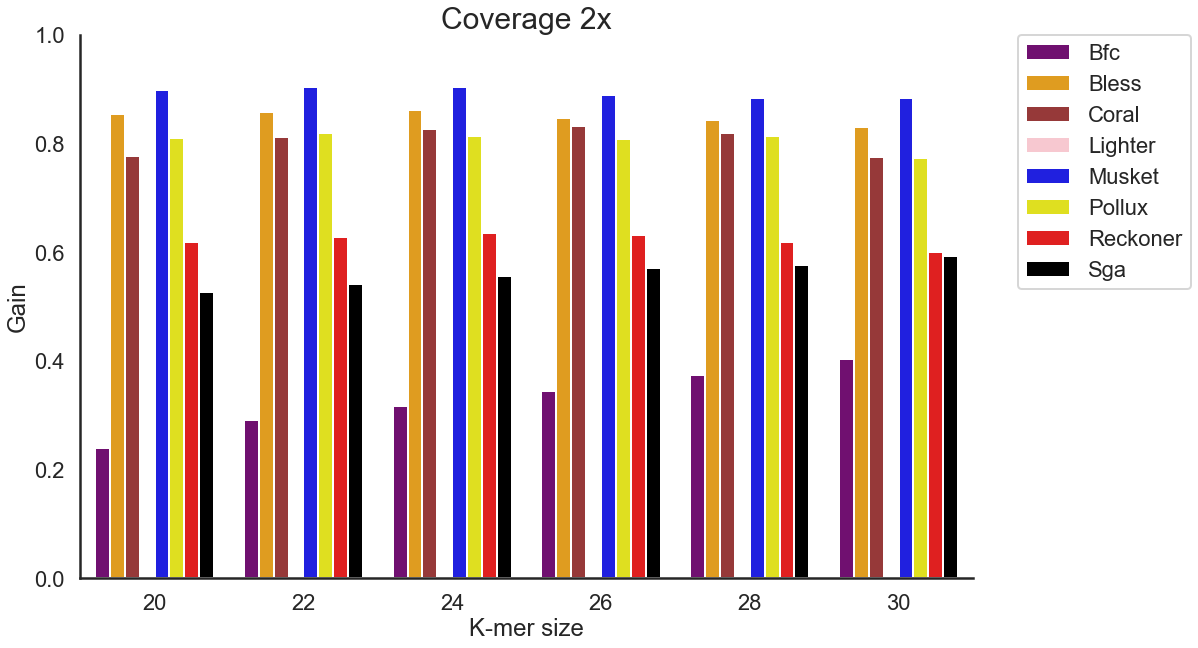

(c)

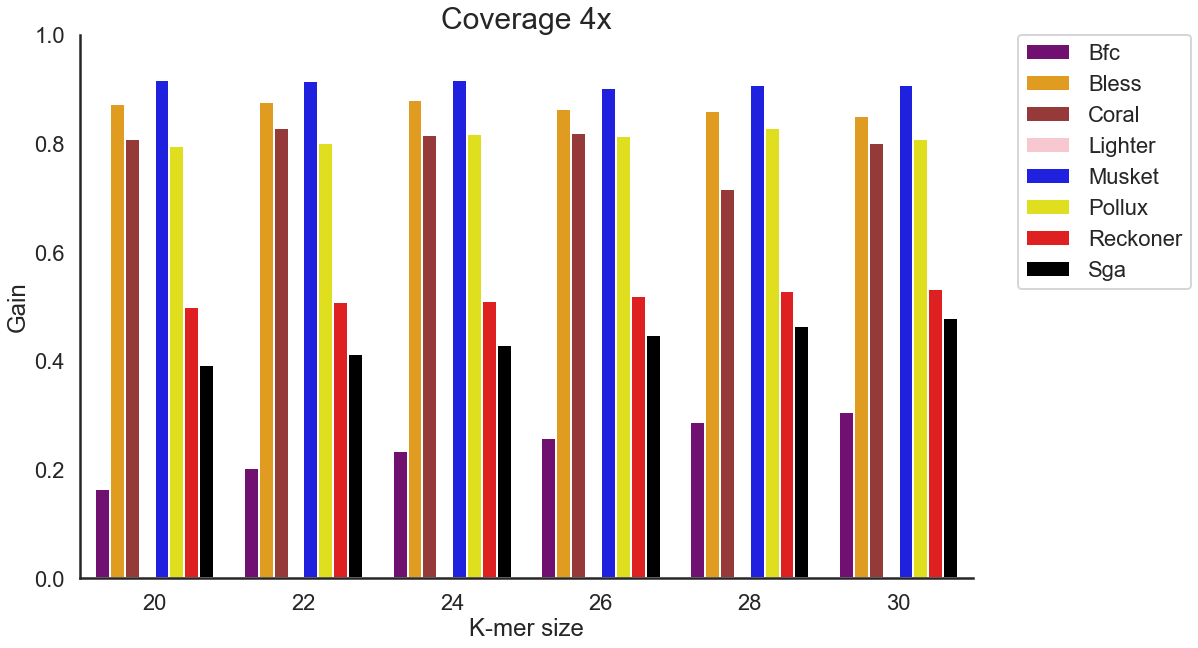

(d)

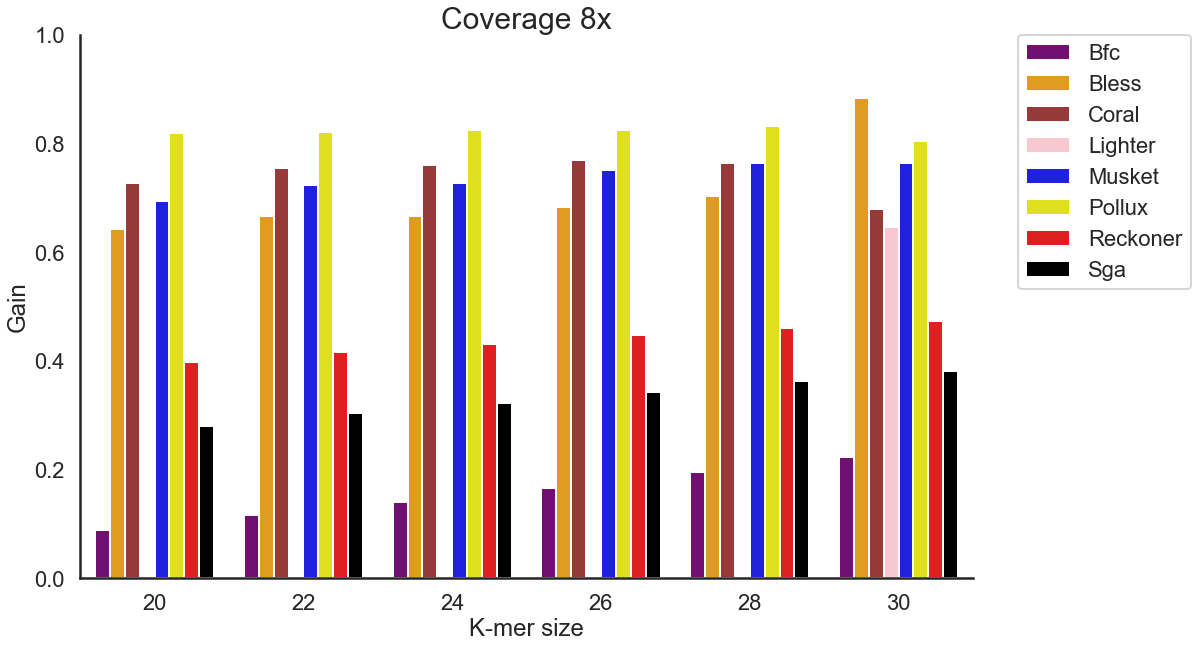

(e)

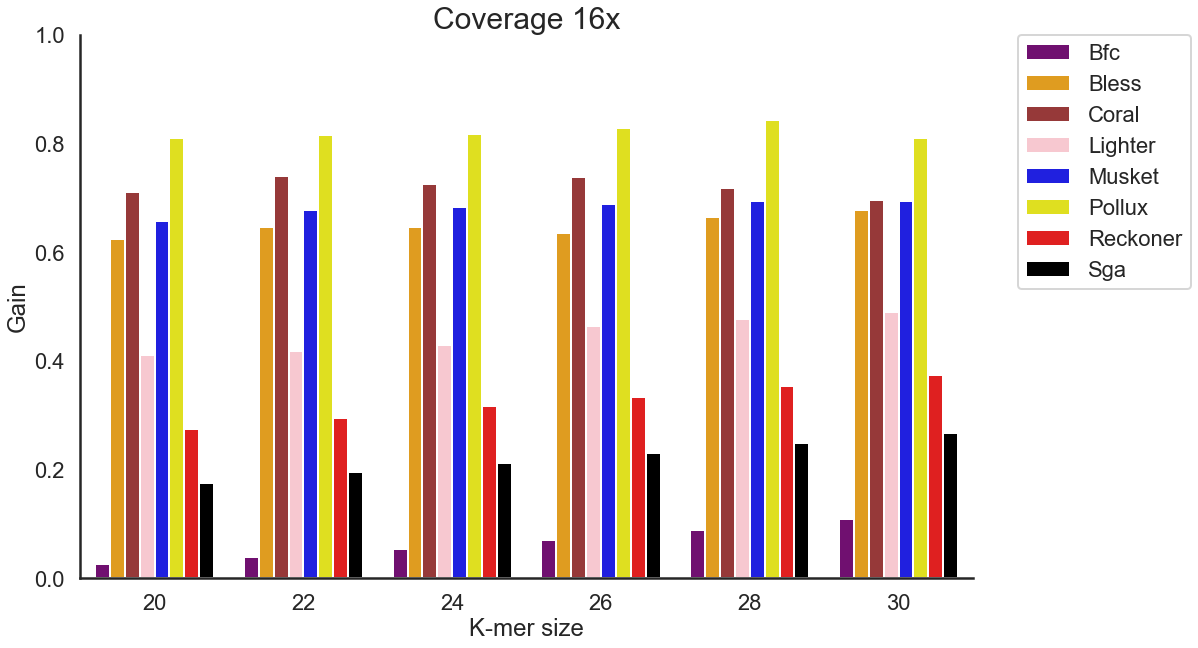

(f)

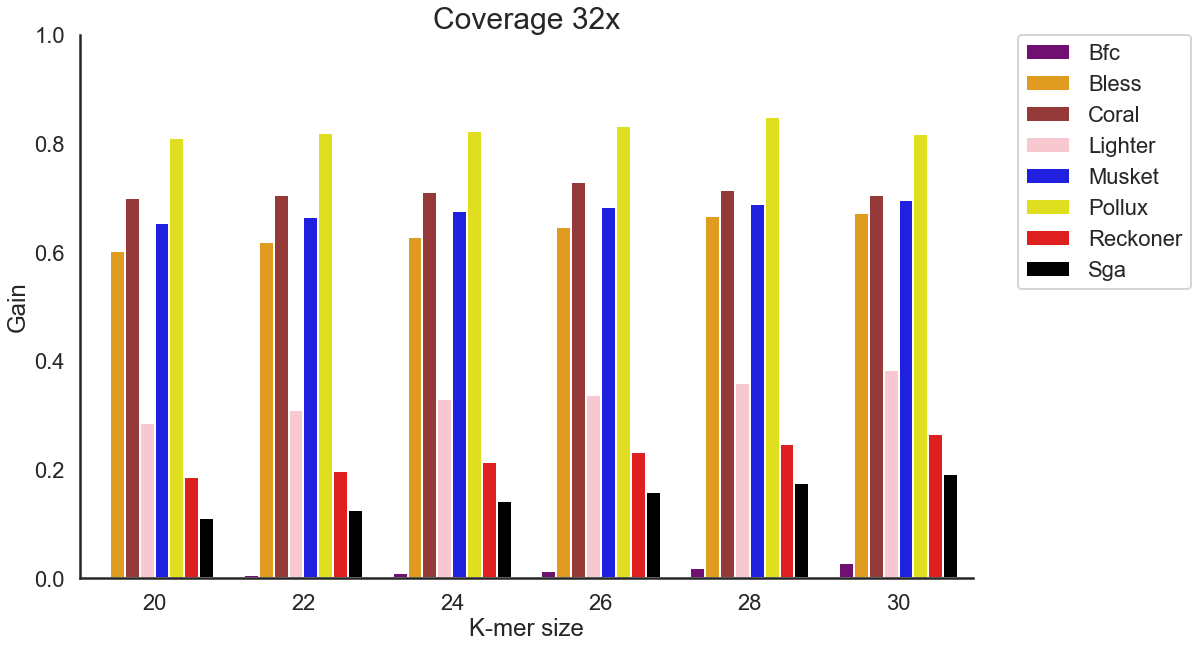

**Figure S12**. The effect of k-mer size on the accuracy of the error correction tools across various coverages settings for TCR-Seq data (D3 dataset). The reads were simulated from TCRA transcripts. (D3 dataset) (a) Coverage of 1x. (b) Coverage of 2x. (c) Coverage of 4x. (d) Coverage of 8x. (e) Coverage of 16x. (f) Coverage of 32x. We have excluded Fiona and Racer, as those tools do not provide options for k-mer size.

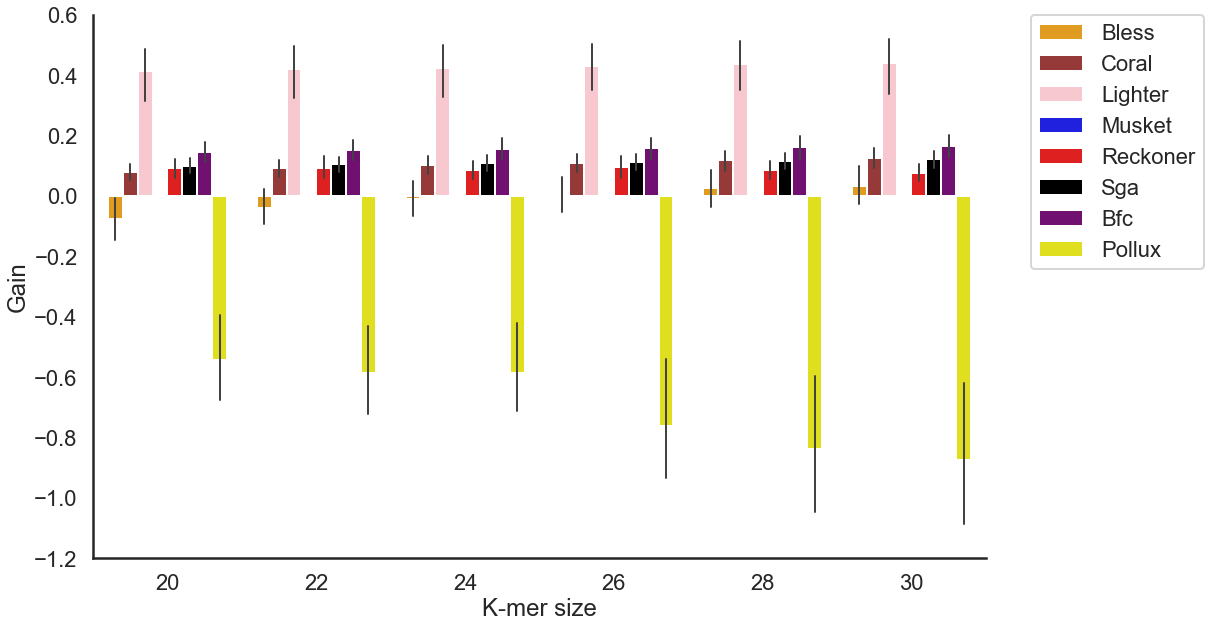

**Figure S13.** The effect of k-mer size on the accuracy of the error correction tools across various coverages settings across 8 real TCR-Seq samples (D2 dataset). TCR-Seq reads were obtained from HIV patients. Error-free reads for gold standard were obtained using UMI-based protocol. We have excluded Fiona and Racer, as those tools do not provide options for k-mer size.

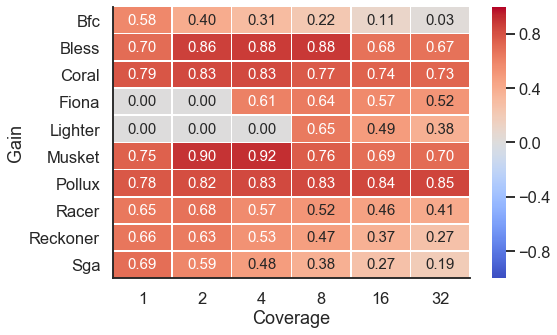

**Figure S14.** Heatmap depicting the gain across various coverage settings of TCR-seq (D3 dataset). Each row corresponds to an error correction tool, and each column corresponds to a dataset with a given coverage.

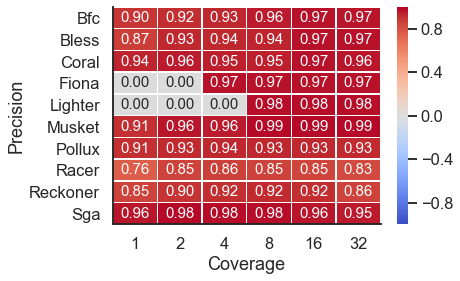

**Figure S15.** Heatmap depicting the precision across various coverage settings of TCR-seq (D3 dataset). Each row corresponds to an error correction tool, and each column corresponds to a dataset with a given coverage.

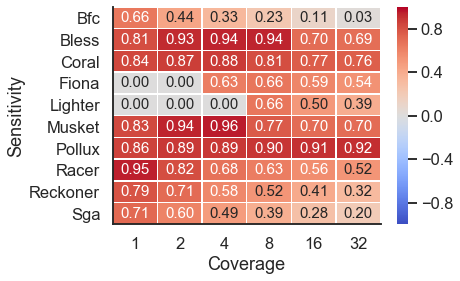

**Figure S16.** Heatmap depicting the sensitivity across various coverage settings of TCR-seq (D3 dataset). Each row corresponds to an error correction tool, and each column corresponds to a dataset with a given coverage.

(a)

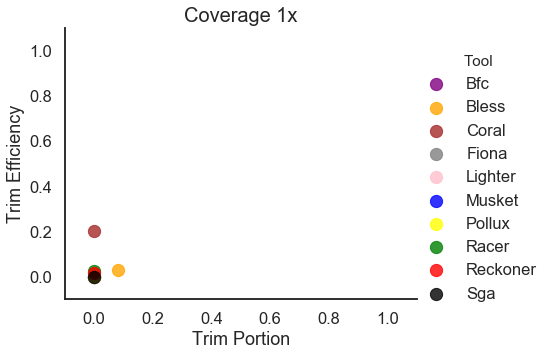

(b)

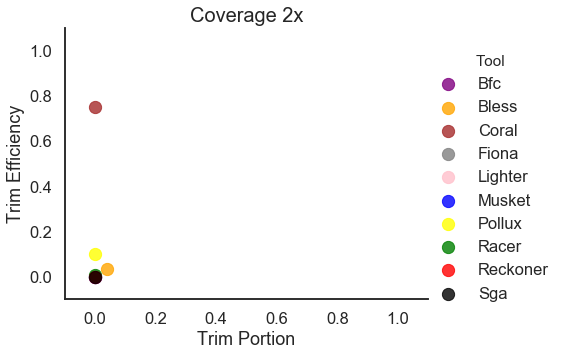

(c)

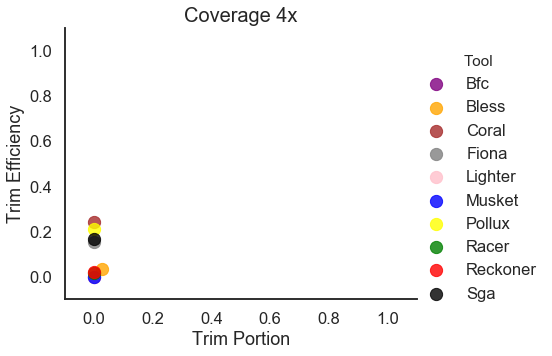

(d)

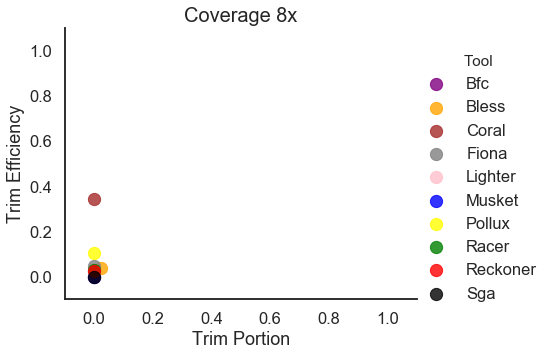

(e)

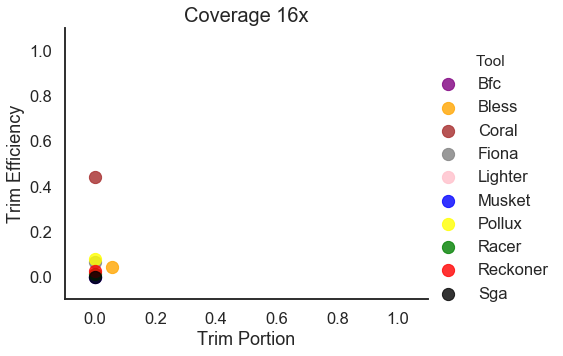

(f)

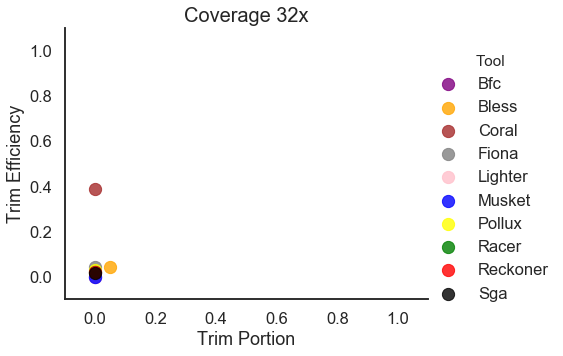

**Figure S17.** Trimming Efficiency vs. Trim Proportion for TCR-Seq simulated data (D3 dataset). The reads were simulated from TCRA transcripts. (a) Coverage of 1x. (b) Coverage of 2x. (c) Coverage of 4x. (d) Coverage of 8x. (e) Coverage of 16x. (f) Coverage of 32x. For each tool, the best k-mer size was selected.

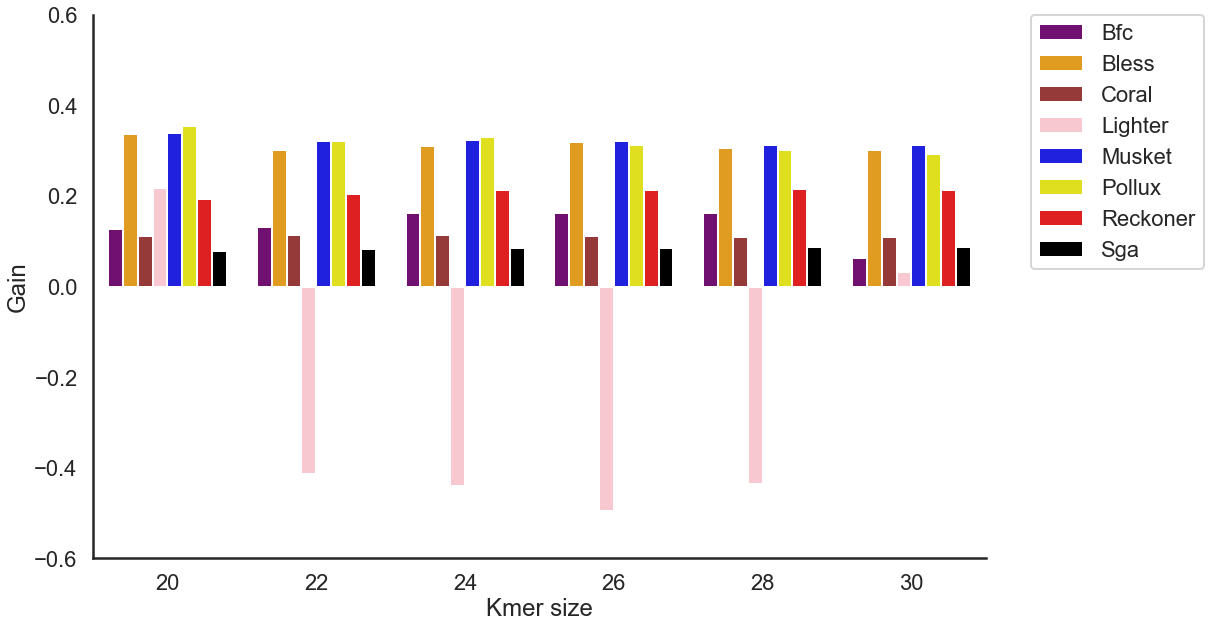

**Figure S18.** The effect of k-mer size on the accuracy of the error correction tools for viral sequencing data (D4 dataset). Error-free reads for gold standard were obtained using UMI-based protocol. We have excluded Fiona and Racer, as those tools do not provide options for k-mer size.

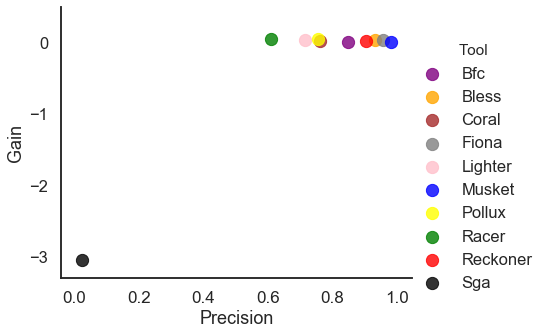

**Figure S19.** Scatter depicting the gain (x-axis) and precision (y-axis) of each tool when applied to D5 HIV mixture dataset. For each tool, the best k-mer size was selected.

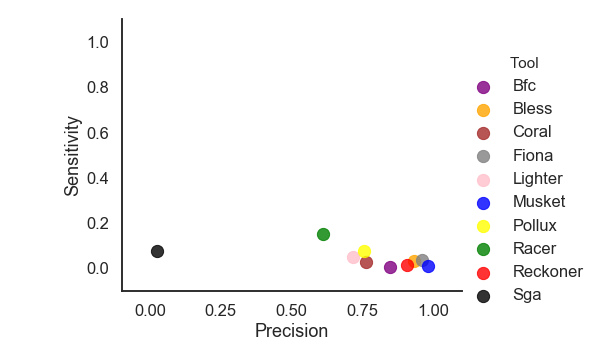

**Figure S20.** Scatter plot depicting the sensitivity (x-axis) and precision (y-axis) of each tool when applied to D5 HIV mixture dataset. For each tool, the best k-mer size was selected.

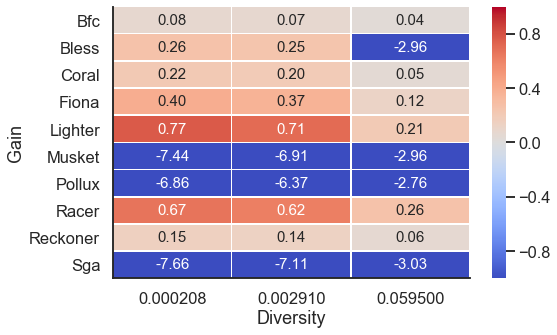

**Figure S21.** Heatmap depicting the gain across D5 HIV mixture dataset for k-mer size set to 20 (with various rates of diversity between haplotypes).

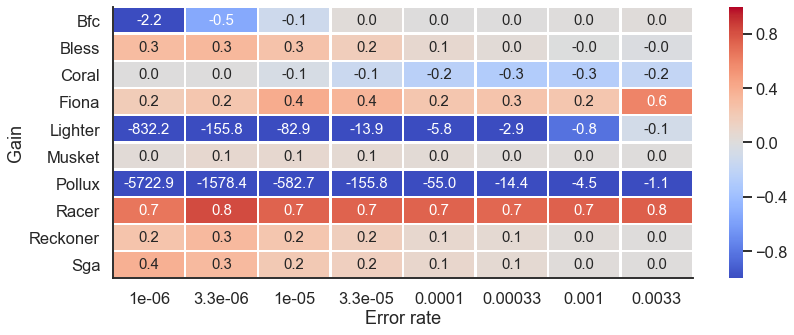

**Figure S22.** Heatmap depicting the gain across D5 HIV mixture dataset (with various error rates). For each tool, the best k-mer size was selected.

**Table S1.** Instructions for running the tools. Datasets D1, D3, D4 and D5 are paired-end files and D2 is single-end files. Input files for executing the tools include the fastq files: input.fastq for single-end files; input_1.fastq and input_2.fastq for paired-end files; merged_input.fastq corresponds to the concatenation of input_1.fastq and input_2.fastq. Other input parameters are: k-mer size (k-mer), genome length (glen), output directory (out_dir), output file (output.fastq and output.fasta).

| **Tool** | **Single-end file** | **Paired-end file** |
| --- | --- | --- |
| BFC | ./bfc -k k-mer input.fastq > output.fastq | ./bfc -k k-mer merged_input.fastq > output.fastq |
| Bless | ./bless -read input.fastq -prefix output.fastq -k-merlength k-mer | ./bless -read1 input_1.fastq -read2 input_2.fastq -prefix output.fastq -k-merlength k-mer |
| Coral | ./coral -fq input.fastq -p 1 -o output.fastq -k k-mer | ./coral -fq merged_input.fastq -p 1 -o output.fastq -k k-mer |
| Fiona | ./fiona -g glen input.fastq output.fasta | ./fiona -g glen merged_input.fastq output.fasta |
| Lighter | ./lighter -r input.fastq -K k-mer glen | ./lighter -r merged_input.fastq -K k-mer glen |
| Musket | ./musket -k k-mer 134217728 -o output.fastq input.fastq | ./musket -k k-mer 134217728 -o output.fastq merged_input.fastq |
| Pollux | ./pollux -i input.fastq -p -o outdir -k k-mer | ./pollux -i input_1.fastq input_2.fastq -p -o outdir -k k-mer |
| Racer | ./racer input_file.fastq output_file.fastq glen | ./racer merged_input_file.fastq output_file.fastq glen |
| Reckoner | ./reckoner -read input.fastq -k-merlength k-mer -prefix outdir -threads 1 | ./reckoner -read merged_input.fastq -k-merlength k-mer -prefix outdir -threads 1 |
| SGA | ./sga preprocess input.fastq -o output.preprocessed.fastq  ./sga index -a ropebwt output.preprocessed.fastq  ./sga correct -k k-mer -o output.fastq output.preprocessed.fastq | ./sga preprocess merged_input.fastq -o output.preprocessed.fastq  ./sga index -a ropebwt output.preprocessed.fastq  ./sga correct -k k-mer -o output.fastq output.preprocessed.fastq |

**Table S2.** Best k-mer size for each tool for Dataset D1.

|  | Bfc | Bless | Coral | Lighter | Musket | Pollux | Reckoner | Sga |
| --- | --- | --- | --- | --- | --- | --- | --- | --- |
| Human Cov 1 | 30 | 30 | 20 | 20 | 28 | 30 | 30 | 30 |
| Human Cov 2 | 30 | 28 | 22 | 20 | 28 | 30 | 30 | 30 |
| Human Cov 4 | 30 | 24 | 26 | 22 | 24 | 30 | 28 | 22 |
| Human Cov 8 | 20 | 30 | 30 | 20 | 28 | 30 | 30 | 28 |
| Human Cov 16 | 24 | 30 | 30 | 26 | 28 | 30 | 30 | 30 |
| Human Cov 32 | 30 | 30 | 30 | 30 | 28 | 30 | 30 | 30 |
| E.coli Cov 1 | 26 | 26 | 20 | 20 | 28 | 30 | 24 | 20 |
| E.coli Cov 2 | 26 | 26 | 20 | 20 | 28 | 24 | 30 | 20 |
| E.coli Cov 4 | 20 | 30 | 20 | 20 | 28 | 20 | 22 | 20 |
| E.coli Cov 8 | 20 | 20 | 20 | 20 | 20 | 20 | 20 | 20 |
| E.coli Cov 16 | 20 | 22 | 20 | 22 | 20 | 22 | 20 | 22 |
| E.coli Cov 32 | 20 | 26 | 20 | 26 | 24 | 26 | 26 | 30 |

**Table S3.** Best k-mer size for each tool for Dataset D2.

|  | Bfc | Bless | Coral | Lighter | Musket | Pollux | Reckoner | Sga |
| --- | --- | --- | --- | --- | --- | --- | --- | --- |
| SRR1543964 | 30 | 24 | 30 | 30 | 20 | 20 | 22 | 30 |
| SRR1543965 | 30 | 30 | 28 | 30 | 20 | 20 | 22 | 30 |
| SRR1543966 | 26 | 30 | 30 | 30 | 20 | 20 | 26 | 30 |
| SRR1543967 | 30 | 30 | 30 | 28 | 20 | 20 | 26 | 30 |
| SRR1543968 | 30 | 28 | 30 | 26 | 20 | 20 | 26 | 30 |
| SRR1543969 | 30 | 30 | 30 | 30 | 20 | 20 | 20 | 30 |
| SRR1543970 | 30 | 28 | 30 | 28 | 20 | 20 | 26 | 30 |
| SRR1543971 | 30 | 30 | 30 | 30 | 20 | 20 | 22 | 30 |

**Table S4.** Best k-mer size for each tool for Dataset D3.

|  | Bfc | Bless | Coral | Lighter | Musket | Pollux | Reckoner | Sga |
| --- | --- | --- | --- | --- | --- | --- | --- | --- |
| TCRA simul.  Cov 1 | 30 | 20 | 26 | 20 | 22 | 22 | 20 | 26 |
| TCRA simul.  Cov 2 | 30 | 24 | 26 | 20 | 24 | 22 | 24 | 30 |
| TCRA simul.  Cov 4 | 30 | 24 | 22 | 20 | 20 | 28 | 30 | 30 |
| TCRA simul.  Cov 8 | 30 | 30 | 26 | 30 | 28 | 28 | 30 | 30 |
| TCRA simul.  Cov 16 | 30 | 30 | 22 | 30 | 28 | 28 | 30 | 30 |
| TCRA simul.  Cov 32 | 30 | 30 | 26 | 30 | 30 | 28 | 30 | 30 |

**Table S5.** Best k-mer size for each tool for Dataset D4.

|  | Bfc | Bless | Coral | Lighter | Musket | Pollux | Reckoner | Sga |
| --- | --- | --- | --- | --- | --- | --- | --- | --- |
| HIV | 28 | 20 | 24 | 20 | 20 | 20 | 28 | 30 |

**Table S6.** Best k-mer size for each tool for Dataset D5.

|  | Bfc | Bless | Coral | Lighter | Musket | Pollux | Reckoner | Sga |
| --- | --- | --- | --- | --- | --- | --- | --- | --- |
| HIV  mixture | 30 | 30 | 30 | 20 | 20 | 20 | 30 | 22 |
| HIV mixture  error rate 0.33% | 30 | 20 | 20 | 28 | 28 | 30 | 30 | 30 |
| HIV mixture  error rate  0.1% | 28 | 26 | 20 | 30 | 28 | 30 | 30 | 30 |
| HIV mixture  error rate  0.033% | 26 | 20 | 20 | 28 | 28 | 30 | 24 | 30 |
| HIV mixture  error rate  0.01% | 26 | 26 | 20 | 30 | 26 | 30 | 20 | 30 |
| HIV mixture  error rate  0.0033% | 26 | 24 | 20 | 28 | 28 | 30 | 30 | 30 |
| HIV mixture  error rate  0.001% | 26 | 24 | 20 | 28 | 26 | 30 | 22 | 28 |
| HIV mixture  error rate  0.00033% | 26 | 24 | 20 | 30 | 22 | 30 | 24 | 26 |
| HIV mixture  error rate  0.0001% | 26 | 30 | 20 | 28 | 20 | 30 | 30 | 30 |
